# DNA-PK Promotes DNA End Resection at DNA Double Strand Breaks in G_0_ cells

**DOI:** 10.1101/2021.10.21.465258

**Authors:** Faith C. Fowler, Bo-Ruei Chen, Nicholas Zolnerowich, Wei Wu, Raphael Pavani, Jacob Paiano, Chelsea Peart, André Nussenzweig, Barry P. Sleckman, Jessica K. Tyler

## Abstract

DNA double-strand break (DSB) repair by homologous recombination is confined to the S and G_2_ phases of the cell cycle partly due to 53BP1 antagonizing DNA end resection in G_1_ phase and non-cycling quiescent (G_0_) cells where DSBs are predominately repaired by non-homologous end joining (NHEJ). Unexpectedly, we uncovered extensive MRE11- and CtIP-dependent DNA end resection at DSBs in G_0_ mammalian cells. A whole genome CRISPR/Cas9 screen revealed the DNA-dependent kinase (DNA-PK) complex as a key factor in promoting DNA end resection in G_0_ cells. In agreement, depletion of FBXL12, which promotes ubiquitylation and removal of the KU70/KU80 subunits of DNA-PK from DSBs, promotes even more extensive resection in G_0_ cells. In contrast, a requirement for DNA-PK in promoting DNA end resection in proliferating cells at the G_1_ or G_2_ phase of the cell cycle was not observed. Our findings establish that DNA-PK uniquely promotes DNA end resection in G_0_, but not in G_1_ or G_2_ phase cells, and has important implications for DNA DSB repair in quiescent cells.

## Introduction

DNA double-strand breaks (DSBs) are particularly deleterious lesions which, if left unrepaired, can lead to cell death, or if repaired aberrantly, can lead to oncogenic chromosomal translocations and deletions (Jackson and Bartek 2009). Eukaryotic cells utilize two main mechanisms of DSB repair: non-homologous end joining (NHEJ), where the broken DNA ends are ligated together with minimal processing of the DNA termini; and homologous recombination (HR), which uses a homologous sequence, usually on a sister chromatid, as a template for accurate DNA repair. Because HR relies on a homologous template for accurate repair, HR is mostly restricted to S and G_2_ phases of the cell cycle when sister chromatids exist. On the other hand, cells can employ NHEJ in any phase of the cell cycle and it is the only option in quiescent (G_0_) cells and G_1_ phase cells (Scully et al. 2019).

Extensive DNA end resection of the broken DNA ends, which generates long tracts of 3’ ssDNA overhangs at DSBs, is a critical step in committing the cell to use HR to repair DSBs. DNA end resection is initiated by nucleases MRE11 and CtIP, and subsequently extended by nucleases including EXO1 and DNA2/BLM (Paull and Gellert 1998; Trujillo et al. 1998; Sartori et al. 2007; Gravel et al. 2008; Mimitou and Symington 2008; Zhu et al. 2008; Bunting et al. 2010). The 3’ ssDNA overhangs are quickly bound by the single-stranded binding protein trimer replication protein A (RPA) to stabilize and protect the ssDNA, and later in repair RPA is replaced by the RAD51 recombinase protein that leads to the homology search to find a homologous template to achieve accurate HR repair (Sugiyama and Kowalczykowski 2002; San Filippo et al. 2008; Wright et al. 2018). NHEJ is initiated by the KU70/KU80 heterodimer binding to broken DNA ends (Zahid et al. 2021). KU70/KU80 recruits the DNA-dependent protein kinase catalytic subunit (DNA-PKcs) which together form a complex called DNA-PK (Gottlieb and Jackson 1993; Hammarsten and Chu 1998). Once the DNA-PK complex is formed, the KU heterodimer translocates inwards along the DNA and DNA-PKcs remains at the DNA ends, undergoing activation via conformational changes mediated by autophosphorylation of the ABCDE cluster (Yaneva et al. 1997; Chen et al. 2021b). Recent cryo-EM structures of DNA-PK also implicate dimerization of DNA-PK as important in recruiting downstream NHEJ factors by bringing broken DNA ends together (Chaplin et al. 2021; Zha et al. 2021). In addition to autophosphorylation, DNA-PKcs phosphorylates members of the NHEJ machinery, including the KU heterodimer, XRCC4, XLF, and Artemis (Bartlett and Lees-Miller 2018).

The critical bifurcation point in the choice to use HR or NHEJ to repair DSBs is the processing of broken DNA ends to form single-stranded 3’ DNA overhangs, which blocks NHEJ and commits the cell to HR (Symington and Gautier 2011). Therefore, DNA end resection is tightly regulated to prevent aberrant DNA end resection in G_0_ and G_1_ phase cells, where NHEJ is the major DSB repair pathway. Several factors have been identified as critical DNA end protection factors that limit resection of DNA DSBs including 53BP1, RIF1, and the Shieldin complex. The proposed mechanism of action of 53BP1 and its downstream effectors include acting as a physical barrier to protect DNA ends from nucleases and promoting DNA polymerase α activity to quickly fill in any resected ends (Dev et al. 2018; Mirman et al. 2018; Noordermeer et al. 2018; Setiaputra and Durocher 2019; Paiano et al. 2021). Additionally, KU70/KU80 has also been shown in budding yeast *Saccharomyces cerevisiae* to inhibit DNA end resection in G_1_ and G_2_ phases of the cell cycle, and in S phase in mammalian cells (Lee et al. 1998; Clerici et al. 2008; Shao et al. 2012).

While nuclease activity is largely limited in G_0_/G_1_ phase cells to prevent aberrant DNA end resection, evidence exists suggesting that nuclease-mediated DNA end processing occurs at some DSBs in G_0_/G_1_. For example, Artemis is required to open hairpin-sealed DNA ends generated during V(D)J recombination in lymphocytes (Menon and Povirk 2016). Additionally, DNA end resection has been observed in G_1_ phase after DNA damage at complex DNA lesions (Averbeck et al. 2014; Biehs et al. 2017), suggesting that DNA end resection is not completely inhibited in the absence of sister chromatids. To investigate what additional factors may regulate DNA end resection in cells lacking sister chromatids, we performed a genome-wide CRISPR/Cas9 screen for genes whose inactivation either increases or decreases RPA bound to chromatin after irradiation (IR) in G_0_-arrested murine cells. We discovered, unexpectedly, that KU70, KU80, and DNA-PKcs promote extensive DNA end resection in G_0_ cells, but not in G_1_ or G_2_ phases of the cell cycle.

## Results

### RPA associates with IR-induced DNA DSBs in G_0_ cells

Murine pre-B cells transformed with Abelson murine leukemia virus (termed abl pre-B cells hereafter) continuously proliferate *in vitro* and can be efficiently arrested in G_0_, also referred to as the quiescent state, upon treatment with the abl kinase inhibitor imatinib (Figure S1A) (Bredemeyer et al. 2006; Chen et al. 2021a). To investigate how DNA end resection is regulated in G_0_ cells, we used a flow cytometric approach to assay RPA bound to chromatin after detergent extraction of soluble RPA, as a proxy for ssDNA generated at DSBs after exposing cells to irradiation (IR) (Forment et al. 2012; Chen et al. 2021a). This assay was performed in abl pre-B cell lines deficient in DNA Ligase IV (*Lig4^-/-^*), to maximize our ability to detect chromatin-bound RPA at DSBs, given that completion of NHEJ is prevented in the absence of DNA Ligase IV. We also performed the analysis in *Lig4^-/-^:53bp1^-/-^* abl pre-B cells which lack the DNA end protection protein 53BP1 and accumulate high levels of RPA on chromatin after IR (Chen et al. 2021a). In agreement with our previous work, we detected a high level of chromatin-bound RPA in G_0_-arrested *Lig4^-/-^:53bp1^-/-^* abl pre-B cells after IR, consistent with the role of 53BP1 in DNA end protection (Figure 1A). Surprisingly, we also observed RPA associated with chromatin after IR of G_0_-arrested *Lig4*^-/-^ abl pre-B cells, although at lower levels than in *Lig4^-/-^:53bp1^-/-^* abl pre-B cells (Figure 1A). Moreover, the increase in IR-induced chromatin-bound RPA does not require DNA Ligase IV deficiency as we were able to observe similar results using the RPA flow cytometric assay in wild-type (WT) abl pre-B cells arrested in G_0_ (Figure S1B). These data indicate that extensive DNA end resection occurs at DSBs in G_0_ cells, despite the presence of the DNA end protection proteins 53BP1 and KU70/KU80.

**Figure 1.**
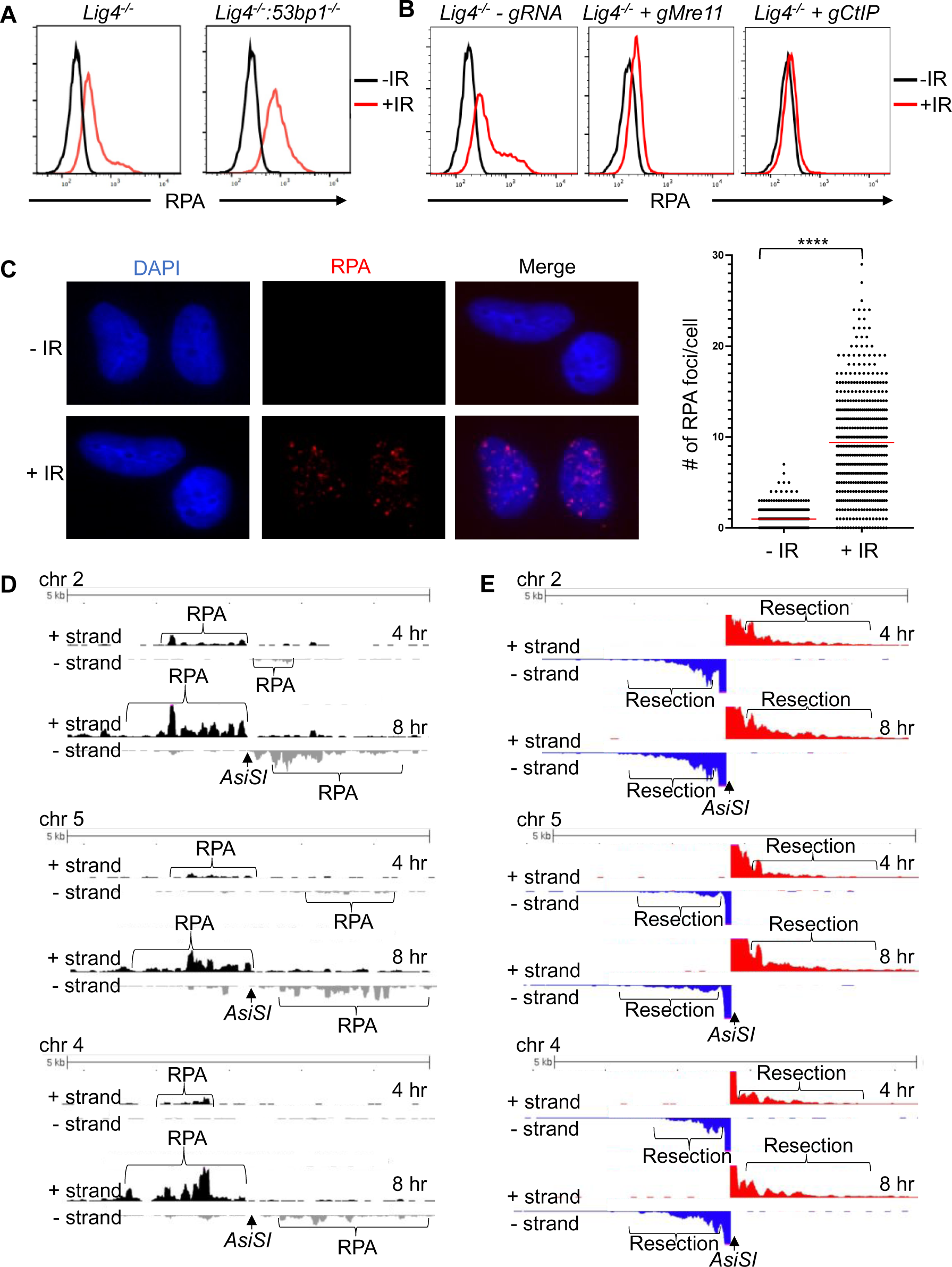
RPA is loaded onto ssDNA after DSBs in G_0_ mammalian cells. (A) Flow cytometric analysis of chromatin-bound RPA in G_0_-arrested *Lig4^-/-^* and *Lig4^-/-^ :53bp1^-/-^* abl pre-B cells before and 3 hours after 20 Gray IR. Representative of three independent experiments. (B) Flow cytometric analysis of chromatin-bound RPA before and 2 hours after 15 Gy IR in G_0_-arrested abl pre-B *Lig4^-/-^* cells (left), *Lig4^-/-^* cells depleted of *Mre11* (middle), and *Lig4^-/-^* cells depleted of *CtIP* (right). Representative of three independent experiments. (C) Representative images and quantification of IR-induced RPA foci from 3 independent experiments in G_0_-arrested MCF10A cells before and 3 hours after 10 Gray IR. n=365 cells in No IR and n=433 cells in IR. Red bars indicate average number of RPA foci in No IR=0.96 and average number of RPA foci in IR=9.4 (****p<0.0001, unpaired t test). (D) RPA ChIP-seq tracks at *AsiSI* DSBs on chromosome 2, 5, and 4 at 4 hours (top) and 8 hours (bottom) after *AsiSI* endonuclease induction in G_0_-arrested *Lig4^-/-^* abl pre-B cells. (E) Representative END-Seq tracks showing resection at *AsiSI* DSBs at chromosome 2, 5, and 4 at 4 hours (top) and 8 hours (bottom) after *AsiSI* induction in G_0_-arrested *Lig4^-/-^* abl pre-B cells. END-seq data is representative from two independent experiments.

To determine whether higher levels of chromatin-bound RPA in irradiated G_0_-arrested *Lig4^-/-^* abl pre-B cells is a result of DNA end resection, we depleted the nucleases that are required for the initiation of DNA end resection during HR in cycling cells. We found that the depletion of CtIP or MRE11 reduced the levels of RPA on chromatin in irradiated G_0_-arrested *Lig4^-/-^* abl pre-B cells (Figure 1B and S1C), indicating that the RPA we observe with our flow cytometric assay after IR is indeed a result of DNA end resection.

To determine whether the DNA end resection that we observed was unique to abl pre-B cells or not, we performed the RPA flow cytometric chromatin association assay in the human breast epithelial cell line MCF10A. We arrested the MCF10A cells in G_0_ by EGF deprivation (Chen et al. 2021a). Similar to *Lig4^-/-^* and WT abl pre-B cells in G_0_, we observed IR-induced chromatin-bound RPA in G_0_ MCF10A cells (Figure S1D), consistent with DNA end resection occurring in these cells at DSBs. RPA binding to ssDNA surrounding DSBs often form distinct nuclear foci that can be easily detected by immunofluorescence staining and microscopy analysis (Golub et al. 1998). Therefore, we performed immunofluorescence staining for RPA in EGF-deprived MCF10A cells. We observed discrete IR-induced RPA foci, consistent with the RPA associated with ssDNA accumulating at DNA damage sites (Figure 1C). Together, these results suggest that broken DNA ends are resected in a CtIP and MRE11-dependent manner, leading to RPA accumulation on ssDNA in G_0_ mammalian cells.

### DNA end resection and RPA loading occurs at site-specific DSBs in G_0_ cells

As irradiation induces DNA base lesions and single-stranded DNA breaks in addition to DSBs, it could potentially complicate our analysis of DNA end processing at regions surrounding DSBs. Therefore, we investigated DSBs at specific locations in the mouse genome upon induction of the *AsiSI* endonuclease. We performed RPA chromatin immunoprecipitation sequencing (RPA ChIP-seq) after induction of *AsiSI* DSBs in G_0_-arrested *Lig4^-/-^* abl pre-B cells. We detected RPA binding adjacent to *AsiSI* DSBs, consistent with ssDNA generated by resection around DNA DSBs (Paiano et al. 2021) (Figure 1D and S1E). Moreover, the association of RPA with chromatin was strand specific around the DSBs, consistent with the 5’-3’ nature of DNA end resection which generates 3’ ssDNA overhangs (Paiano et al. 2021) (Figure 1D). To determine the extent of DNA end processing in G_0_ cells, we performed END-seq (Canela et al. 2016; Wong et al. 2021) to directly measure DNA end resection at nucleotide resolution at *AsiSI*-induced DSBs. Using END-seq, we detected extensive DNA end resection in G_0_-arrested *Lig4^-/-^* abl pre-B cells at 4 and 8 hours after *AsiSI* DSB induction (Figure 1E). Together, these data indicate that in G_0_-arrested cells, DNA ends are resected at DSBs induced by IR or site-specific endonucleases, generating ssDNA that is bound by RPA.

### A CRISPR/Cas9 screen identifies the DNA-PK complex as promoting DNA end resection in G_0_ cells

To identify factors that influence DNA end resection in G_0_ cells, we performed a genome-wide CRISPR/Cas9 screen in G_0_-arrested *Lig4^-/-^* abl pre-B cells 2 hours after irradiation to identify factors that either promote or impair DNA end resection (Figure 2A).

**Figure 2.**
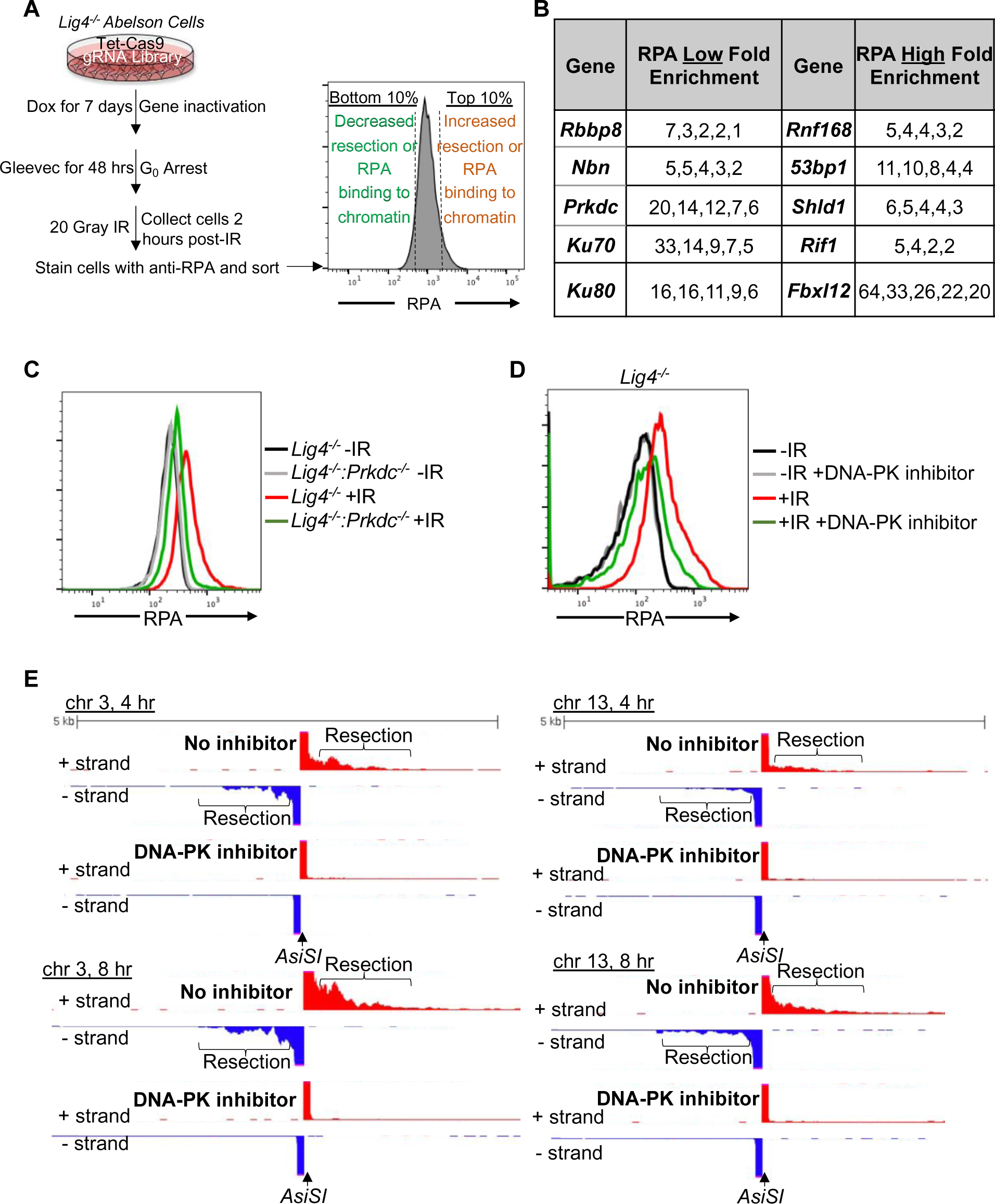
A genome-wide gRNA screen identifies DNA-PK as a factor that promotes DNA end resection in G_0_. (A) Schematic of genome-wide guide RNA screen for factors promoting (low RPA) or inhibiting (high RPA) chromatin-bound RPA loading 2 hours after 20 Gray IR in G_0_-arrested *Lig4^-/-^* abl pre-B cells. (B) Fold enrichment of selected guide RNAs in low RPA high RPA populations. (C) Flow cytometric analysis of chromatin-bound RPA in G_0_-arrested *Lig4^-/-^* and *Lig4^-/-^:Prkdc^-/-^* abl pre-B cells before and 3 hours after 15 Gray IR. Data is representative of three independent experiments in two different cell lines. (D) Flow cytometric analysis of chromatin-bound RPA in G_0_-arrested *Lig4^-/-^* abl pre-B cells with and without 10 μm NU7441 (DNA-PKcs inhibitor) pre-treatment 1 hour before 20 Gray IR. Data is representative of three independent experiments in two different cell lines. (E) Representative END-seq tracks at chromosome 3 (left) and chromosome 13 (right) in G_0_-arrested *Lig4^-/-^* abl pre-B cells 4 hours (top) and 8 hours (bottom) after *AsiSI* DSB induction, with and without 10 μm NU7441 treatment.

We isolated the 10% of cells with the lowest RPA (low RPA) and the 10% cells with the highest RPA (high RPA) staining intensity using our RPA flow cytometric assay followed by flow assisted cell sorting. We then amplified the guide RNAs (gRNAs) in these populations of cells and determined their frequencies using high throughput sequencing. gRNAs enriched in the low RPA staining population correspond to genes encoding proteins that normally promote DNA end resection, while gRNAs enriched in the high RPA population correspond to genes encoding proteins that normally impair resection. In this screen we identified several gRNAs enriched in the low RPA staining population to *Rbbp8* which encodes the nuclease CtIP, and *Nbn,* which encodes the NBN subunit of the MRE11-RAD50-NBN (MRN) complex, consistent with their established roles in promoting DNA end resection (Figure 2B). Unexpectedly, we also found gRNAs of *Ku70*, *Ku80*, and *Prkdc* (the gene encoding DNA-PKcs) highly enriched in our low RPA population (Figure 2B). This suggested that DNA-PK may promote DNA end resection in G_0_ cells, contrary to the established role of these factors in preventing DNA end resection in other phases of the cell cycle.

To validate the screen and determine if DNA-PK is required for DNA end resection, we generated *Lig4^-/-^:Prkdc^-/-^* abl pre-B cells that do not express DNA-PKcs by CRISPR/Cas9-mediated gene inactivation (Figure S2A). G_0_-arrested *Lig4^-/-^:Prkdc^-/-^* abl pre-B cells had lower levels of chromatin-bound RPA after IR compared to *Lig4^-/-^* abl pre- B cells (Figure 2C). DNA-PKcs and Ataxia-telangiectasia mutated (ATM) are two major serine/threonine kinases that are activated in response to DNA DSBs and share some overlapping functions due to similar substrate specificity (Blackford and Jackson 2017). Because DNA-PKcs but not ATM was identified in our screen, we wanted to determine if the pro-resection activity in G_0_-arrested cells is unique to DNA-PKcs or also shared by ATM. We treated G_0_-arrested *Lig4^-/-^* abl pre-B cells with the ATM inhibitor KU55933 or the DNA-PKcs inhibitor NU7441 before IR and performed flow cytometric analysis of IR-induced chromatin-bound RPA. In contrast to the consistent reduction in the levels of chromatin-bound RPA observed in G_0_-arrested *Lig4^-/-^* abl pre-B cells treated with DNA-PKcs inhibitor, ATM inhibition did not have a detectable effect on the levels of IR-induced binding of RPA in G_0_-arrested *Lig4^-/-^* abl pre-B cells (Figure 2D and S2B). The role of DNA-PK in promoting DNA end resection in G_0_ is not limited to murine abl pre-B cells as we also observed a reduced number of IR-induced RPA foci in G_0_-arrested human MCF10A cells upon inhibition of DNA-PKcs (Figure S2C). These results indicate that DNA-PKcs activity, but not ATM, uniquely promotes resection and RPA binding to damaged chromatin after IR in G_0_ cells.

To directly observe if DNA-PKcs influenced DNA end resection at DSBs, we performed nucleotide resolution END-seq on G_0_-arrested *Lig4^-/-^* abl pre-B cells with and without DNA-PKcs inhibitor treatment before the induction of *AsiSI* DSBs. Consistent with our RPA flow cytometric assay results, DNA-PKcs inhibitor-treated G_0_-arrested *Lig4-/-* abl pre-B cells showed greatly reduced END-Seq signals distal to DSBs, consistent with limited DNA end processing when DNA-PK is inactivated (Figure 2E and S2D). These results demonstrate that DNA-PK activity promotes DNA end resection of DSBs in G_0_ mammalian cells.

### FBXL12 inhibits KU70/KU80-dependent DNA end resection in G_0_ cells

Given that DNA-PKcs promotes DNA end resection in G_0_ cells (Figure 2C, 2D, 2E), and that *Ku70* and *Ku80* were enriched in the low RPA loading population of cells in the CRISPR/Cas9 screen (Figure 2B), we determined whether KU70/KU80 may also promote resection in G_0_ cells. We generated *Lig4^-/-^:Ku70^-/-^* abl pre-B cells and measured DNA end resection using our RPA flow cytometric approach. Consistent with our observations in DNA-PK inhibitor-treated G_0_-arrested *Lig4^-/-^* abl pre-B cells and *Lig4^-/-^:Prkdc^-/-^* abl pre-B cells, the level of chromatin-bound RPA after IR was greatly reduced in G_0_-arrested *Lig4^-/-^:Ku70^-/-^* abl pre-B cells compared to *Lig4^-/-^* abl pre-B cells (Figure 3A and S3A). As such, the entire DNA-PK complex is required for DNA end resection in G_0_ cells.

**Figure 3.**
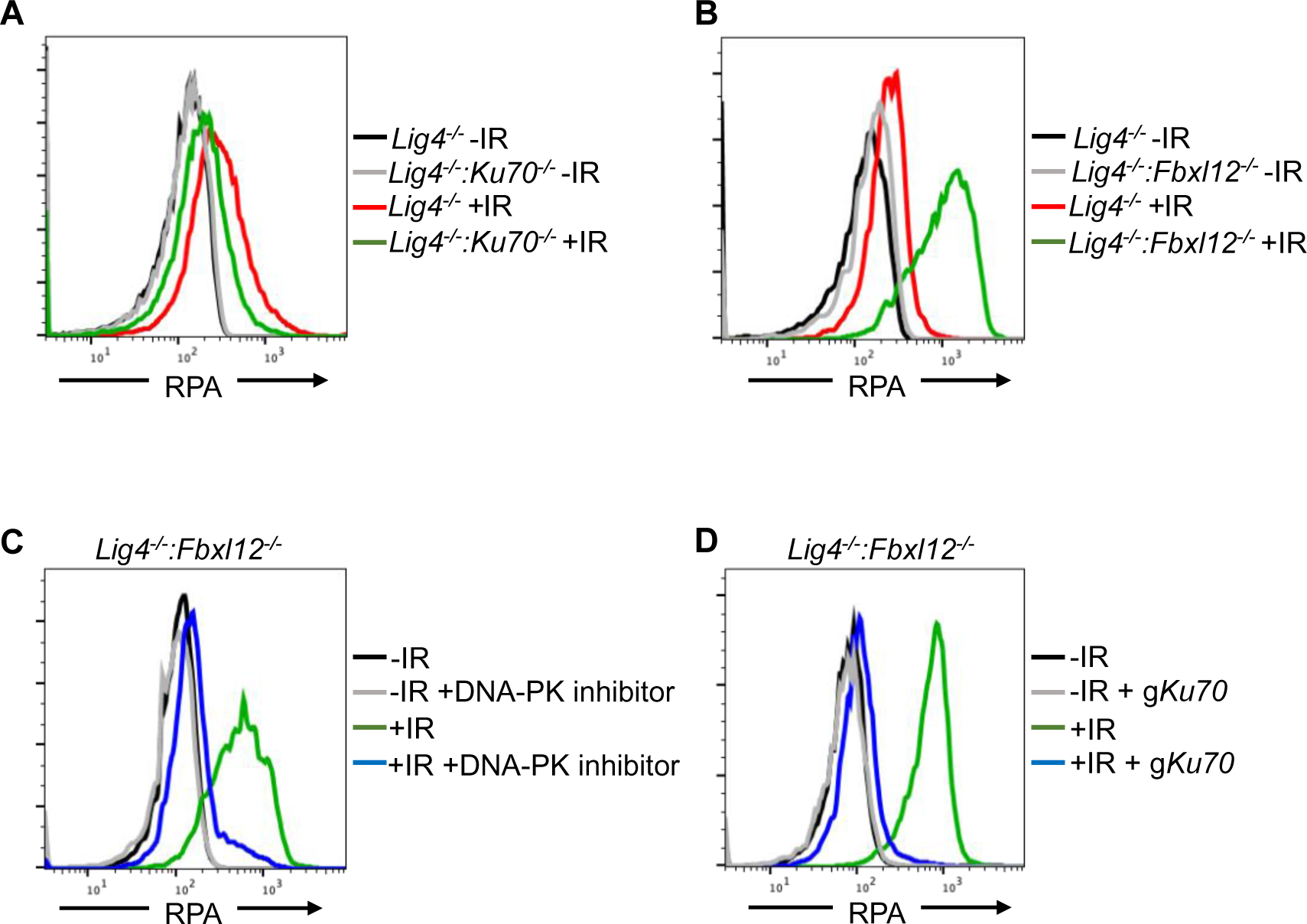
FBXL12 inhibits KU70/KU80-promoted DNA end resection. (A) Flow cytometric analysis of chromatin-bound RPA in G_0_-arrested *Lig4-/-* abl pre-B cells vs *Lig4-/-:Ku70-/-* abl pre-B cells before and 3 hours after 20 Gray IR. Data is representative of three independent experiments in two different cell lines. (B) As in A, in G_0_-arrested *Lig4-/-* and *Lig4-/-:Fbxl12-/-* abl pre-B cells. Data is representative of three independent experiments in at least two different cell lines. (C). Flow cytometric analysis of chromatin-bound RPA in G_0_-arrested *Lig4-/-:Fbxl12-/-* abl pre-B cells with and without 10 μm NU7441 treatment, before and 3 hours after 20 Gray IR. Data is representative of three independent experiments in at least two different cell lines (D) Flow cytometric analysis of chromatin-bound RPA in G_0_-arrested *Lig4-/-:Fbxl12-/-* abl pre-B cells before and after KU70 knockout, before and 3 hours after 15 Gray IR. Data is representative of three independent experiments.

KU70/KU80 is removed from DSBs via ubiquitylation, which has been shown to be mediated by E3 ligases including RNF138, RNF8, RNF126, and the SCF^Fbxl12^ complex (Postow et al. 2008; Feng and Chen 2012; Postow and Funabiki 2013; Ismail et al. 2015; Ishida et al. 2017). In agreement, gRNAs of *Fbxl12*, which encodes the substrate recognition subunit FBXL12 of the SCF^Fbxl12^ E3 ubiquitin ligase complex, were highly enriched in our screen in the high RPA staining cell population (Figure 2B), consistent with the idea that the persistent presence of KU70/KU80 at DSBs in cells lacking FBXL12 would lead to persistent DNA end resection. Indeed, we observed that in G_0_-arrested *Lig4^-/-^:Fbxl12^-/-^* abl pre-B cells, the level of IR-induced chromatin-bound RPA increased compared to *Lig4^-/-^* abl pre-B cells (Figure 3B and Fig S3B). Given the role of FBXL12 on limiting the levels of KU70/KU80 at broken DNA ends, we tested whether the increased DNA end resection phenotype in *Lig4^-/-^:Fbxl12^-/-^* abl pre-B cells depended on DNA-PK activity or the presence of the KU70/KU80 complex. Indeed, inhibition of DNA-PK with NU7441 (Figure 3C) and depletion of KU70 (Figure 3D and Fig S3C) in G_0_-arrested *Lig4^-/-^:Fbxl12^-/-^* abl pre-B cells prevented excessive accumulation of RPA on chromatin after IR. Our results suggest that the ability of DNA-PK to promote DNA end resection in G_0_ cells is regulated through maintaining proper levels of KU70/KU80 at DNA DSBs by the SCF^Fbxl12^ E3 ubiquitylation complex.

### DNA-PK uniquely promotes DNA end resection exclusively in G_0_ cells

KU70/KU80 have been shown to prevent DNA end resection in G_1_ and G_2_ phases in budding yeast and in S phase in mammalian cells but has not been examined in G_0_ cells (Lee et al. 1998; Clerici et al. 2008; Shao et al. 2012). Thus, we set out to determine whether DNA-PK-dependent DNA end resection is limited to G_0_ or can occur in other phases of the cell cycle. To this end, we compared the levels of IR-induced chromatin bound RPA in *Lig4^-/-^*, *Lig4^-/-^:Prkdc^-/-^* and *Lig4^-/-^:Ku70^-/-^* abl pre-B cells arrested in G_0_ by imatinib, arrested in G_2_ by the CDK1 inhibitor RO3306, and in G_1_ phase (cells with 2N DNA) in a proliferating population. In contrast to G_0_ cells, loss of DNA-PKcs (*Lig4^-/-^:Prkdc^-/-^*) did not reduce the levels of IR-induced chromatin-bound RPA in G_2_-arrested or cycling G_1_ phase cells (Figure 4A and S4A). Similar results were obtained when analyzing *Lig4^-/-^:Ku70^-/-^* abl pre-B cells (Figure 4B). The unique function of DNA-PK activity in promoting DNA end resection in G_0_-arrested cells was confirmed with END-seq analysis of *AsiSI*-induced DSBs in *Lig4^-/-^* abl pre-B cells arrested in G_0_ or G_2_ and treated with or without DNA-PKcs inhibitor. Whereas G_0_-arrested *Lig4^-/-^* abl pre-B cells treated with DNA-PKcs inhibitor exhibited significantly reduced END-seq signals in regions distal to the DSBs, the same treatment had little effect in cells arrested in G_2_ phase of the cell cycle (Figure 4C and S4B). These results suggest that DNA-PK distinctly promotes DNA end resection at DSBs in G_0_ but not in other cell cycle phases.

**Figure 4.**
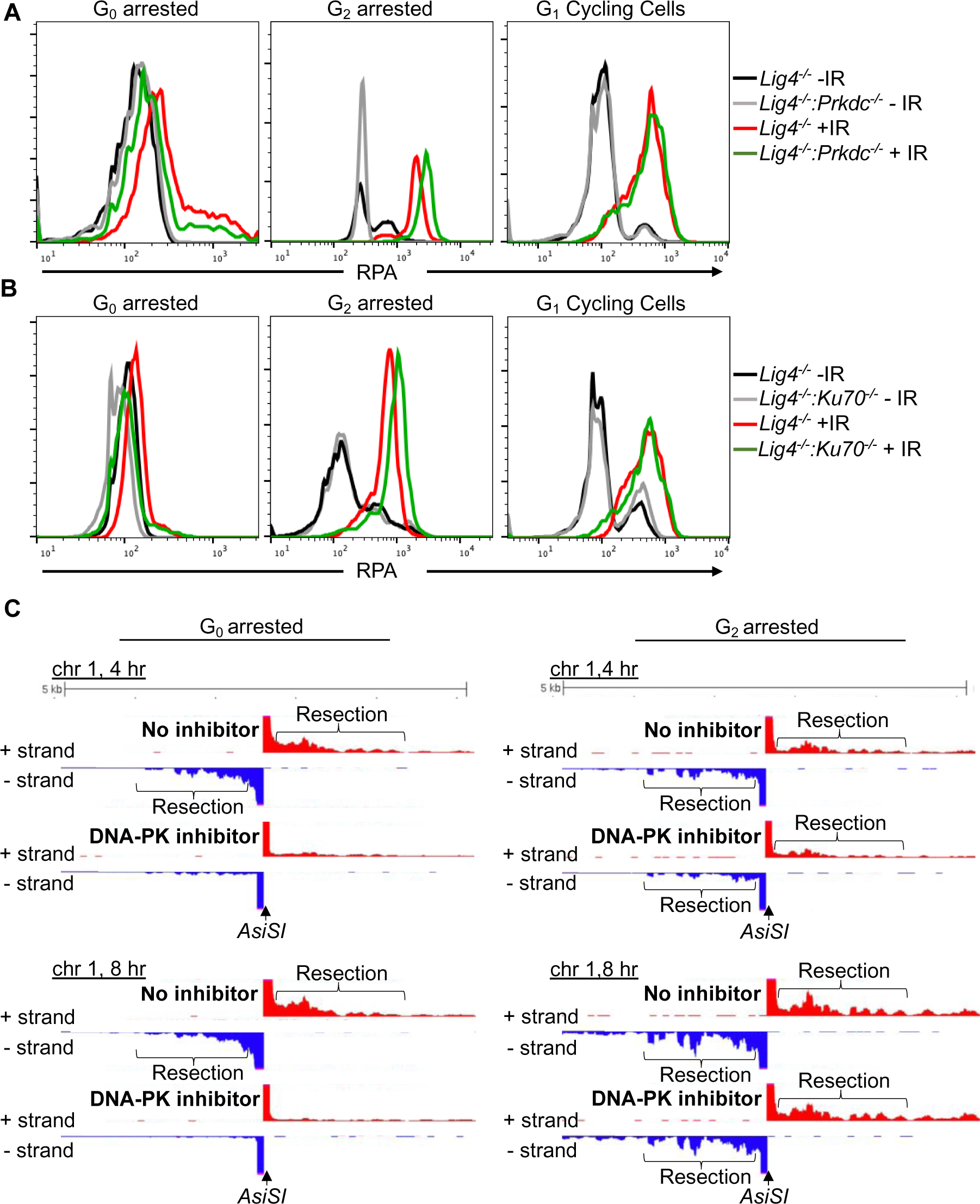
DNA-PK mediates DNA end resection in G_0_ but not G_1_ or G_2_. (A) Flow cytometric analysis of chromatin-bound RPA in *Lig4^-/-^* and *Lig4^-/-^:Prkdc^-/-^* abl pre- B cells arrested in G_0_ (left), arrested in G_2_ by 10 μm RO-3306 treatment for 16 hours and gated on 4N (middle), and G_1_ cells gated on 2N DNA content in cycling cells (right), before and 3 hours after 20 Gray IR. Data is representative of three independent experiments in at least two different cell lines. (B) As in A in *Lig4^-/-^* and *Lig4^-/-^:Ku70^-/-^* abl pre-B cells. (C) Representative END-seq tracks in G_0_ (left) and G_2_-arrested (right, by 10 μm RO-3306 treatment for 16 hours) *Lig4^-/-^* abl pre-B cells, with and without 10 μm NU7441 treatment on chromosome 1, 4 hours (top) and 8 hours (bottom) after *AsiSI* endonuclease induction.

## Discussion

DNA end resection is one of the key events that determines whether cells utilize NHEJ, HR, or other repair pathways utilizing homologous sequences. During G_0_ and G_1_ phase of the cell cycle, NHEJ is the predominant DSB repair pathway and DNA end resection is largely limited compared to other phases of the cell cycle. However, in this study we revealed that DNA end resection dependent on CtIP and MRE11, which are required for resection in S and G_2_ phases of the cell cycle, occurs at DSBs in G_0_ mammalian cells (Figures 1B). In addition to CtIP and MRE11, we identified additional factors that promote resection in G_0_ cells as components of the DNA-PK complex, including KU70, KU80 and DNA-PKcs, in a genome-wide CRISPR/Cas9 screen and showed that the kinase activity of DNA-PK is critical as resection of DSBs diminishes upon DNA-PKcs inhibitor treatment (Figures 2 and 3). Interestingly, we also found in our genome-wide CRISPR/Cas9 screen that inactivating FBXL12, the substrate recognition subunit of the SCF^FBXL12^ E3 ubiquitin ligase complex, promotes extensive resection of DNA ends in G_0_ cells (Figure 3B). As the SCF^FBXL12^ E3 ubiquitin is thought to limit the abundance of the KU70/KU80 heterodimer (Postow and Funabiki 2013), our data are in line with the notion that loss of FBXL12 results in aberrant accumulation of KU70/KU80 at DSBs, and consequently elevated or prolonged activation of DNA-PK at DSBs which promotes resection in G_0_ cells (Figure 5).

**Figure 5.**
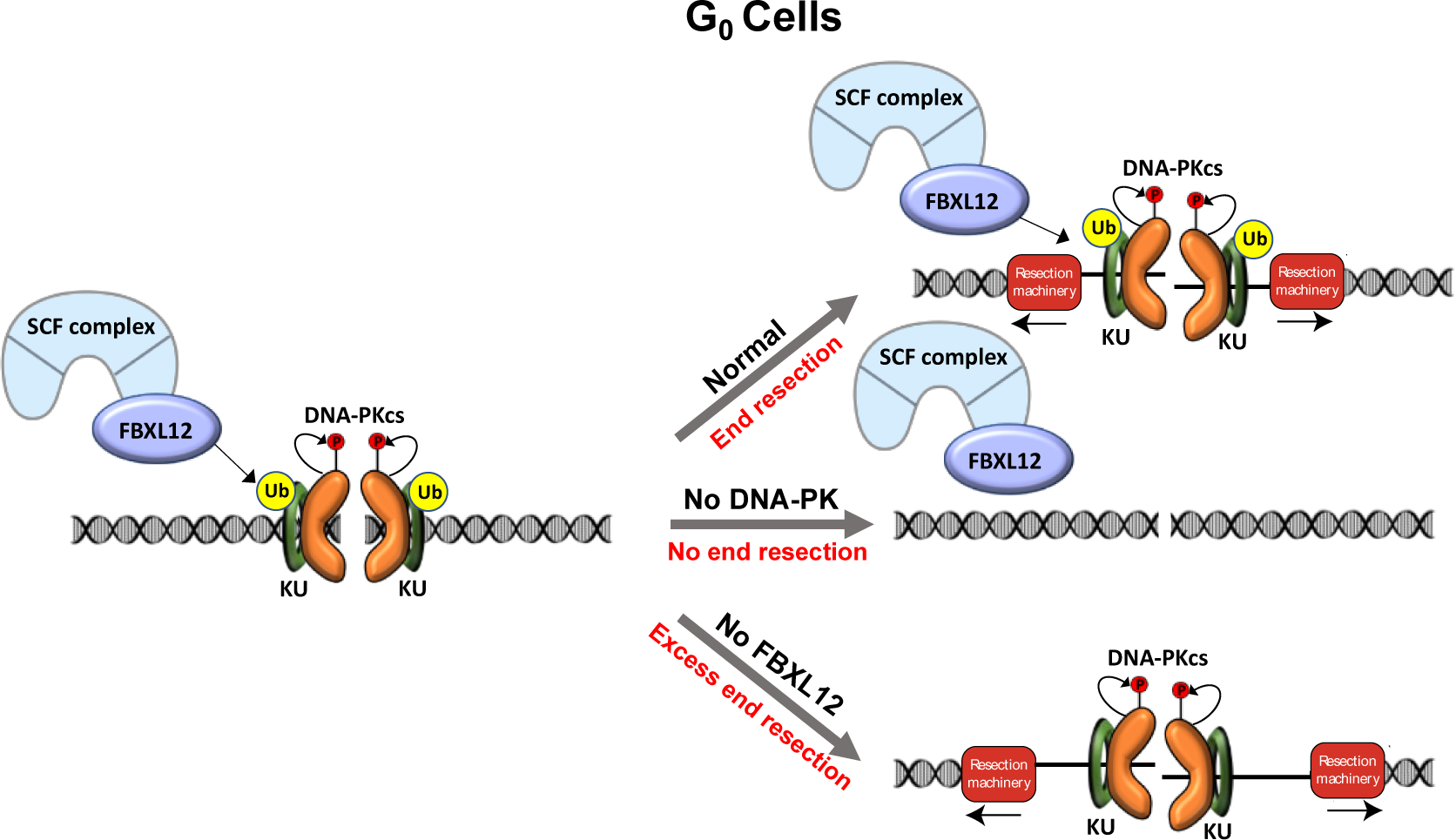
Model of DNA-PK-mediated DNA end resection in G_0_ cells. Normally in G_0_ at DSBs, the DNA-PK complex promotes DNA end resection. This resection is counteracted by FBXL12. Without DNA-PK, there is no DNA end resection in G_0_. Without FBXL12, DNA-PK persists at DSBs which leads to more extensive DNA end resection.

Why would resection occur in G_0_ cells? Chemical modifications or secondary structures at DSBs have been identified as requiring DNA end processing to create a more accessible repair environment, which could presumably be the case at DSBs in G_0_ cells (Weinfeld and Soderlind 1991). For example, Artemis is an endo and exonuclease which is activated by DNA-PKcs and uses its nuclease activity to open DNA hairpins at coding ends, which is required for V(D)J recombination, and cleaves 3’ ssDNA overhangs during NHEJ (Ma et al. 2002; Ma et al. 2005). Though Artemis was not identified in our screen as having a role in G_0_ DSB repair, it serves as an example of nuclease activity being critical for DSB repair outside of HR. Interestingly, DNA end resection has a role in recruiting anti-resection factors to limit extensive DNA end resection. The SHLD2 component of Shieldin binds ssDNA, suppresses RAD51 loading, and ultimately recruits 53BP1 to DSBs (Noordermeer et al. 2018). HELB, a 5’-3’ DNA helicase, binds to RPA and limits EXO1 and BLM-DNA2-mediated DNA end resection (Tkáč et al. 2016). Limited DNA end resection in G_0_ cells could be important in preventing extensive DNA end resection. Altogether, we propose that DNA end resection in G_0_ cells is likely not resulting in aberrant HR but may be required to create more accessible DNA ends and/or to recruit anti-resection factors.

Studies investigating the role of KU70/KU80 during DSB repair have found that KU70/KU80 protects DSBs from nuclease activity. For example, at HO endonuclease breaks in budding yeast, deletion of KU70/KU80 leads to ssDNA accumulation in G_1_ cells and increased MRE11 recruitment to DSBs compared to wild-type cells (Lee et al. 1998; Clerici et al. 2008). Also in budding yeast, at inducible I-SceI DSBs, deletion of KU70 results in increased RFA1 foci formation in G_1_, but deletion of NHEJ factor DNA Ligase IV leads to no defect in RFA1 foci formation compared to wild-type cells, indicating that KU70 itself, not NHEJ, is a barrier to DNA end resection (Barlow et al. 2008). In mammalian cells, complementation of KU70/KU80 knockout cells with a *M. tuberculosis* KU homolog persistently bound to DSBs in S phase results in reduced RPA and RAD51 foci formation after IR (Shao et al. 2012). Contrary to these roles for KU70/KU80 in protecting DNA ends from nucleolytic attack, we found that in G_0_ cells, KU70/KU80 promotes DNA end resection (Figure 3A and 4B). We hypothesize that KU70/KU80 promotes resection through recruitment and activation of DNA-PKcs at DSBs (Gottlieb and Jackson 1993), as we also found that DNA-PKcs inhibition and genetic deletion of *Prkdc* leads to more RPA on chromatin after IR and more DNA end resection in G_0_ cells (Figure 2C-E, S2C, S2D, 4A, 4C). It is important to note that most studies establishing the role of KU70/KU80 in protecting DNA ends were performed in *S. cerevisiae* which do not have a homolog to DNA-PKcs. Therefore, we hypothesize that the function of DNA-PK promoting DNA end resection in G_0_ cells may not be evolutionarily conserved. Moreover, previous studies in *S. cerevisiae* and mammalian cells establishing DNA-PK as a pro-NHEJ complex did not analyze G_0_ cells. We found that DNA-PK does not promote DNA end resection in G_1_ or G_2_ phase cells, only in G_0_-arrested cells, indicating that DNA-PK-dependent DNA end resection is unique to G_0_, but is not contradictory to its anti-resection function in G_1_ or G_2_ phase cells (Figure 4). In G_0_ cells, KU70/KU80 could protect some DNA ends, but after recruitment and activation of DNA-PKcs, the net effect is DNA end resection. Additional studies may elucidate how the balance between DNA end protection and DNA end resection is regulated in G_0_.

ATM and DNA-PK have been shown to have some overlapping functions in DNA damage response and repair, including phosphorylation of H2A.X in response to IR and signal join formation during V(D)J recombination (Stiff et al. 2004; Zha et al. 2011). However, we find that this is not the case during DNA end resection in G_0_ cells as DNA-PK promotes resection in G_0_ cells, but ATM does not have a detectable impact (Figure S2B). ATM has been implicated in promoting HR repair by phosphorylating CtIP and promoting KU removal from DSBs, as well as phosphorylating DNA-PKcs at single-ended DSBs to remove it from these breaks that require DNA end resection (Wang et al. 2013; Britton et al. 2020). DNA-PKcs autophosphorylation promotes HR by removing it from DSBs to allow nuclease access, but is typically associated with promoting NHEJ by phosphorylating Artemis, XRCC4, and XLF (Zhou and Paull 2013; Bartlett and Lees-Miller 2018) So while ATM often promotes DNA end resection and HR, it appears that DNA-PKcs could be acting in place of ATM to promote DNA end resection in G_0_ cells. It is additionally possible that DNA-PKcs phosphorylates a unique substrate(s) in G_0_ cells that promotes DNA end resection.

In summary, we provide here evidence that DNA-PK promotes DNA end resection uniquely in G_0_ cells, and that this DNA end resection is counteracted by FBXL12. We speculate that some aspects of DSB repair in G_0_ function differently than DSB repair in cycling cells, and future studies may reveal the mechanism and utility of these key differences.

## Acknowledgements

The authors thank Chitra Mohan for designing the model graphic and Yinan Wang for performing the bioinformatics for the high throughput screen. We thank the Weill Cornell Flow Cytometry Core for flow cytometry and thank the Weill Cornell Epigenomics Core for providing advice and performing the sequencing for the high throughput screen. JKT is supported by NIH R35 GM139816 and R01 CA95641. BPS is supported by NIH R01 AI047829 and R01 AI074953. FCF is supported by NIH F31 CA239442. JKT and BPS were also supported by the Starr Cancer Consortium and Emerson Collective Cancer Research Fund.

---

**Figure S1.**
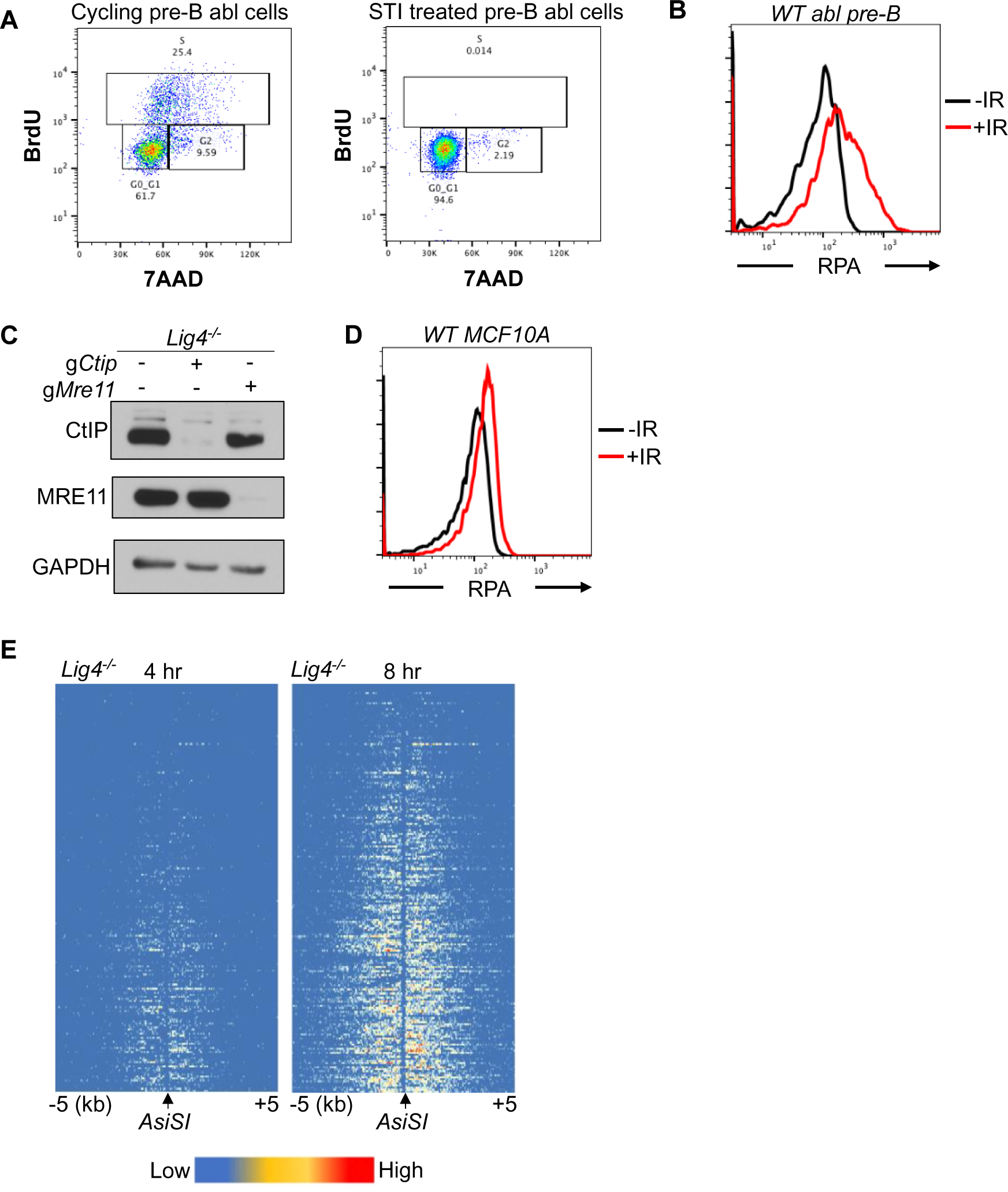
RPA is loaded onto ssDNA after DSBs in G_0_ mammalian cells. (A) Flow cytometric analysis of cycling and STI treated *Lig4^-/-^* abl pre-B cells for BrdU content (y-axis) and DNA content (7AAD, x-axis). (B) Flow cytometric analysis of chromatin-bound RPA in wild-type G_0_-arrested abl pre-B cells before and 3 hours after 20 Gray IR. (C) Western blot of bulk CtIP and MRE11 knockout in *Lig4^-/-^* abl pre-B cells (D) Flow cytometric analysis of chromatin-bound RPA loading in wild-type G_0_-arrested MCF10A cells before and 3 hours after 20 Gray IR. (E) Heat maps of RPA ChIP-seq results at top 200 *AsiSI* sites in G_0_-arrested *Lig4^-/-^* abl pre-B cells 4 hours (left) and 8 hours (right) after *AsiSI*-endonuclease induction.

**Figure S2.**
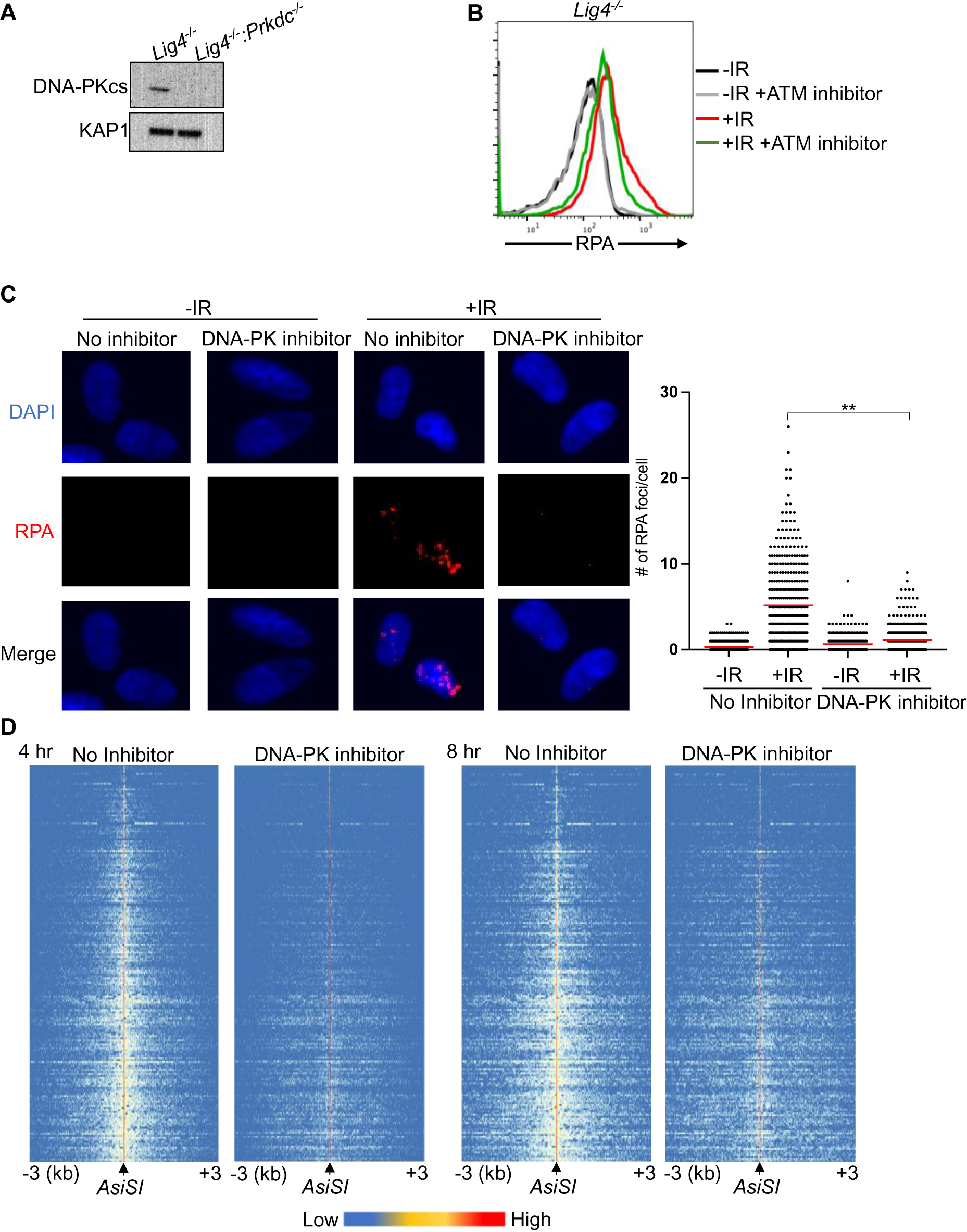
A genome-wide gRNA screen identifies DNA-PK as a factor that promotes DNA end resection in G_0_. (A) Western blot analysis of DNA-PKcs protein in *Lig4^-/-^* and *Lig4^-/-^:Prkdc^-/-^* abl pre-B cells. (B) Flow cytometric analysis of chromatin-bound RPA in G_0_-arrested abl pre-B cells with and without 1 hour pre-treatment with 15 μm KU-55933 (ATM inhibitor), before and 3 hours after 20 Gray IR. (C) Representative images and quantitation from 3 independent experiments of IR-induced RPA foci in G_0_-arrested MCF10A cells with and without 10 μm NU7441, before and 3 hours after 10 Gray IR. For No Inhibitor -IR condition, n=426, average number of RPA foci=0.34. For No Inhibitor +IR condition, n=389, average number of RPA foci=5.2. For DNA-PK inhibitor treated -IR condition, n=266, average number of RPA foci=0.66. For DNA-PK inhibitor treated +IR condition, n=441, average number of RPA foci=1.13. Red bar indicates mean number of RPA foci (**p=0.003). (D) Heat maps of END-seq at top 200 *AsiSI* DSBs with and without 10 μm NU7441 treatment in G_0_-arrested *Lig4^-/-^* abl pre-B cells 4 hours (left) and 8 hours (right) after *AsiSI* DSB induction.

**Figure S3.**
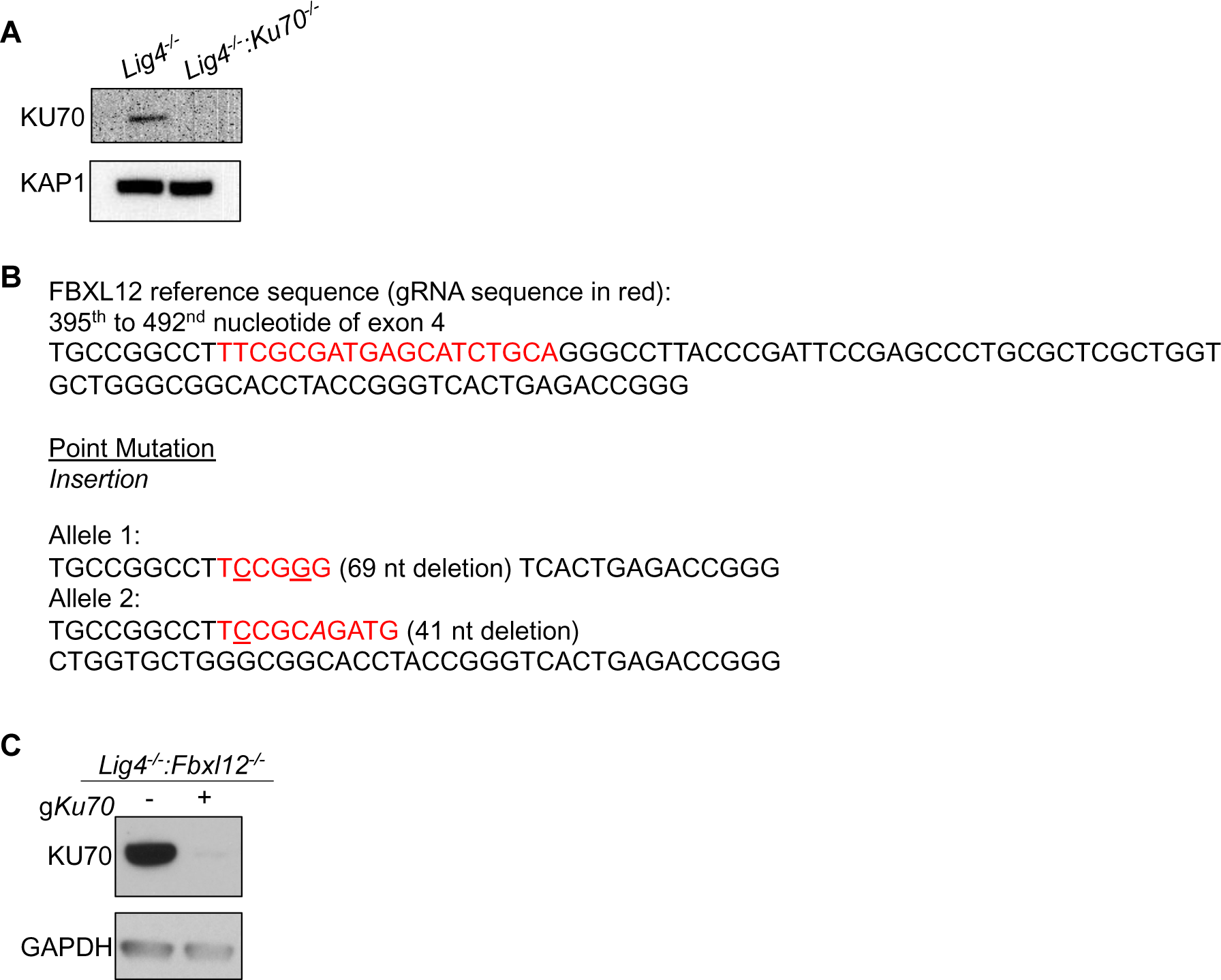
FBXL12 inhibits KU70/KU80-promoted DNA end resection. (A) Western blot analysis of KU70 protein in *Lig4-/-* and *Lig4-/-:Ku70-/-* abl pre-B cells. (B) Gene sequence of *Lig4^-/-^:Fbxl12^-/-^* abl pre-B cell clones indicating deletions. (C) Western blot analysis of KU70 protein in *Lig4^-/-^:Fbxl12^-/-^* abl pre-B cells.

**Figure S4.**
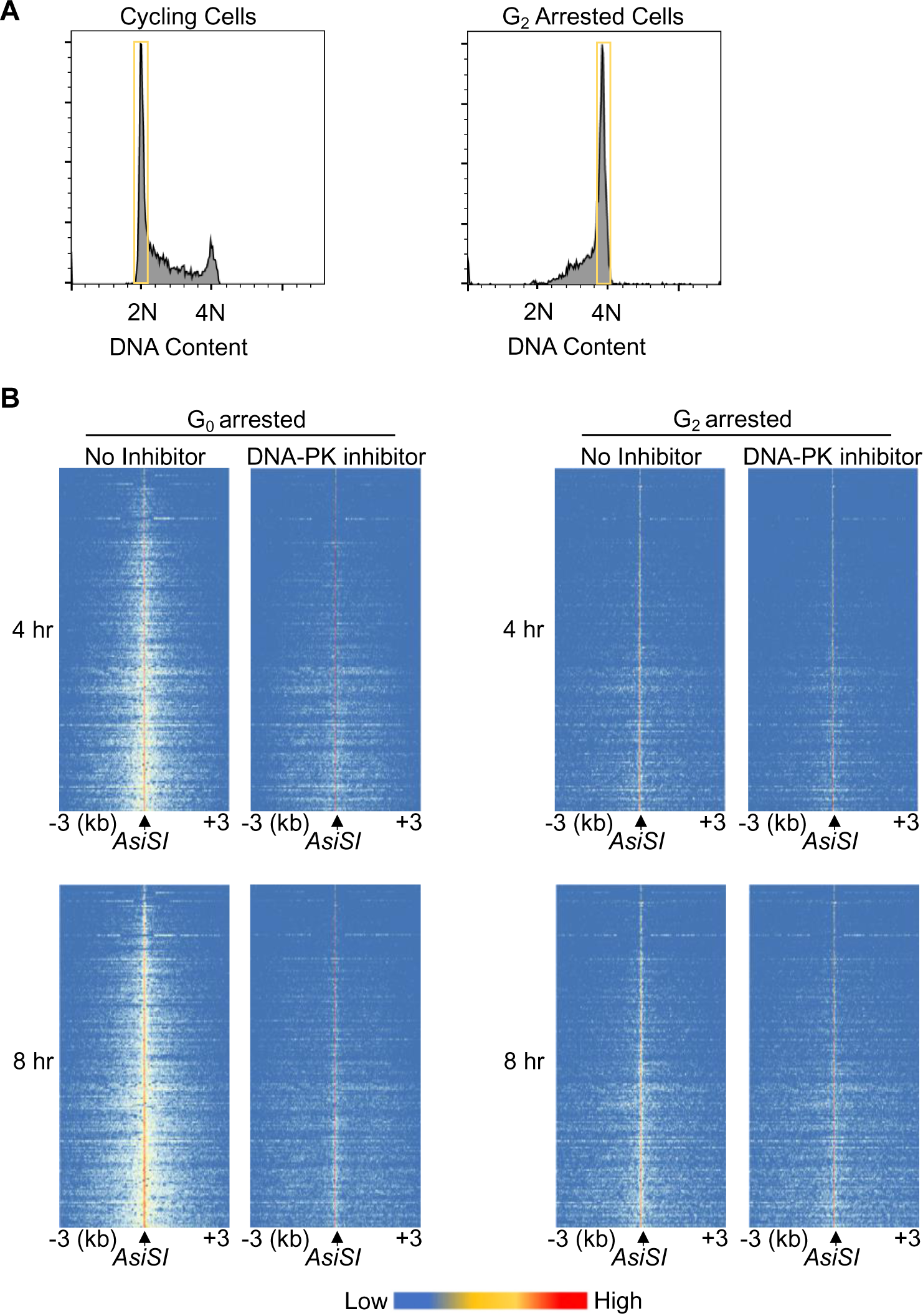
DNA-PK mediates DNA end resection in G_0_ but not G_1_ or G_2_. (A) Flow cytometric analysis of DNA content (7AAD) in RO-3306 treated (right) and cycling cells (left) showing gating used for RPA flow cytometry analysis. (B) Heat maps of END-seq at top 200 *AsiSI* DSBs in G_0_-arrested *Lig4^-/-^* abl pre-B (left) and G_2_-arrested *Lig4^-/-^* abl pre-B (right) 4 hours (top) and 8 hours (bottom) after *AsiSI* DSB induction, with and without 10 μm NU7441 treatment.

## Materials and Methods

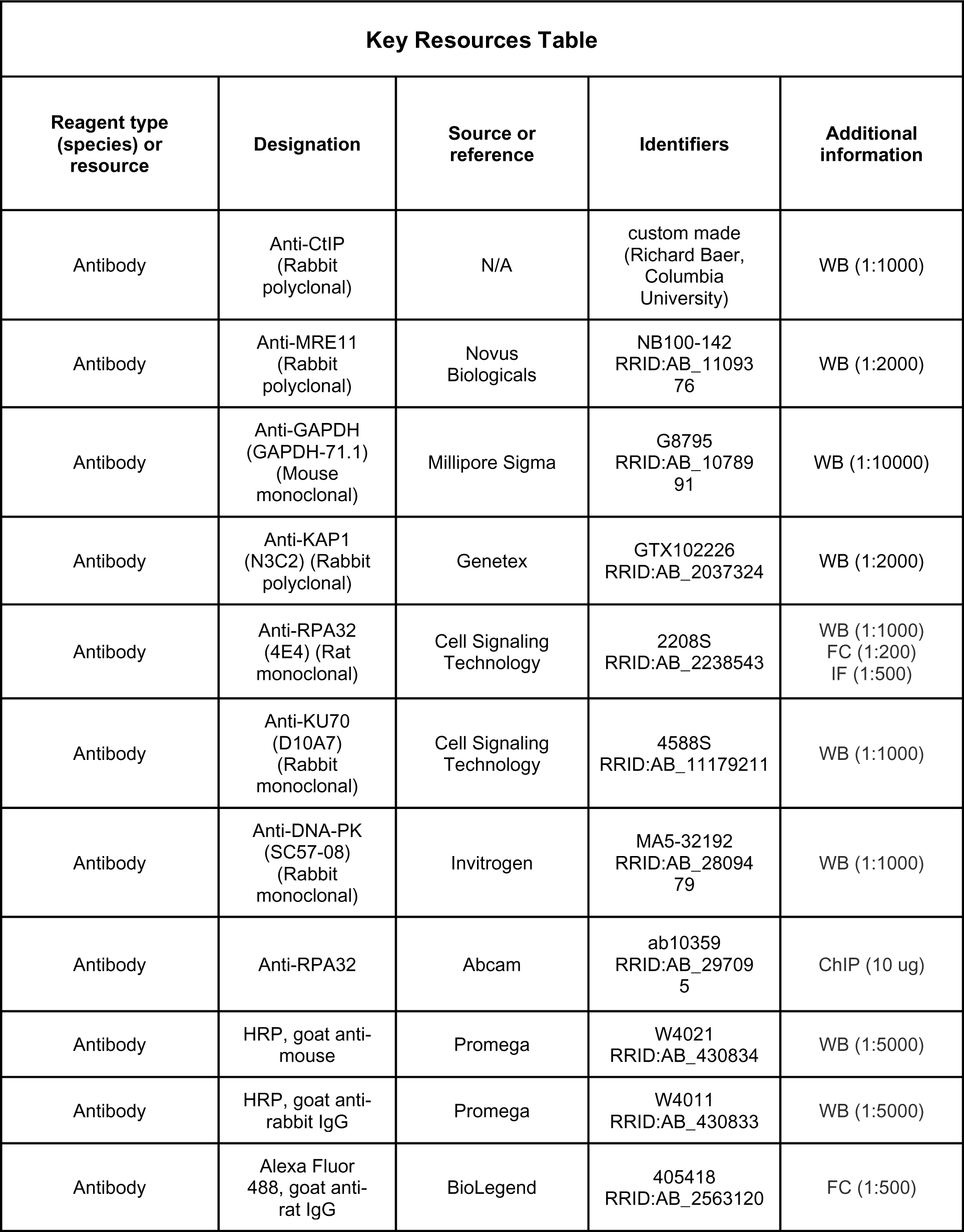
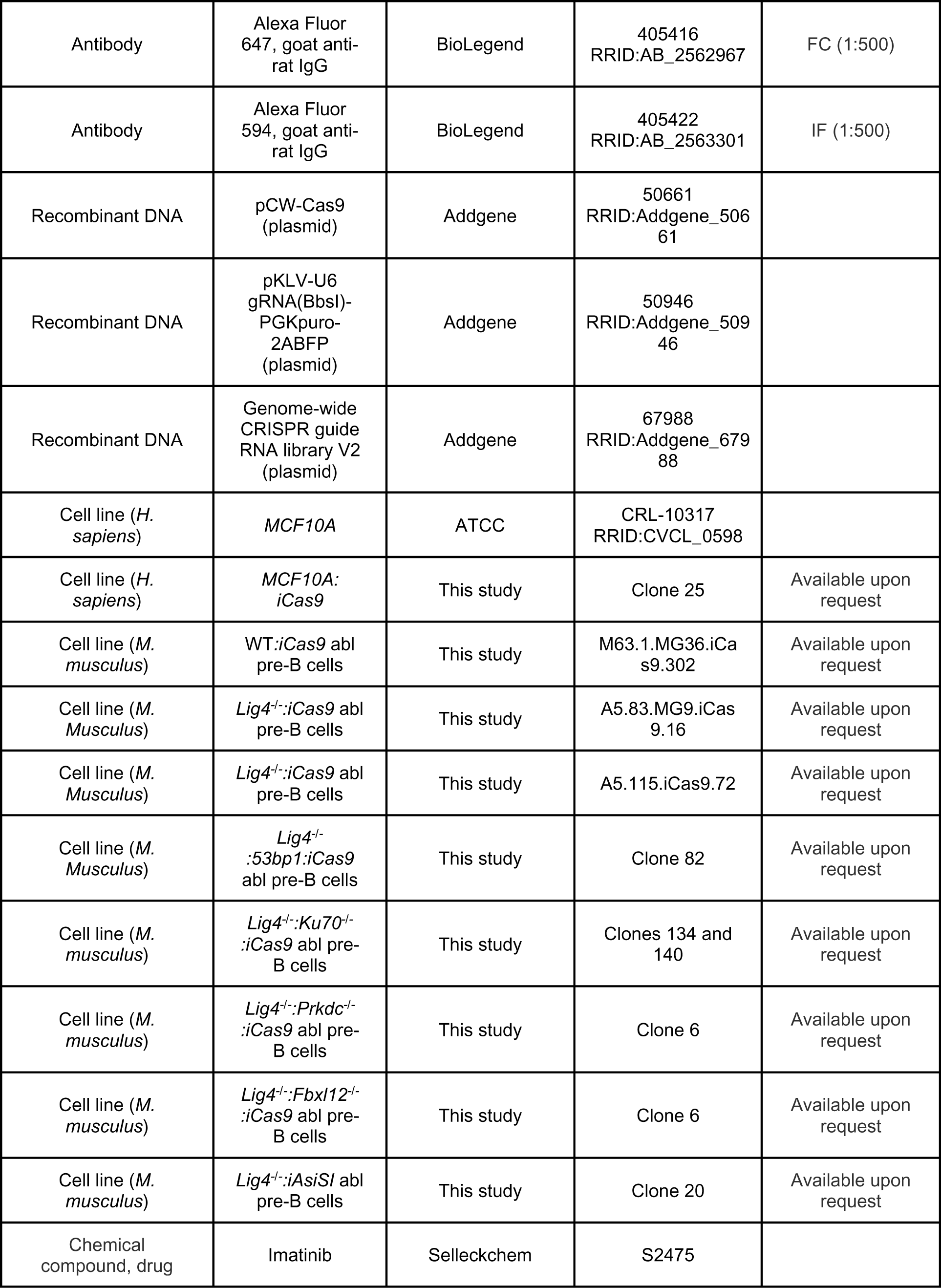
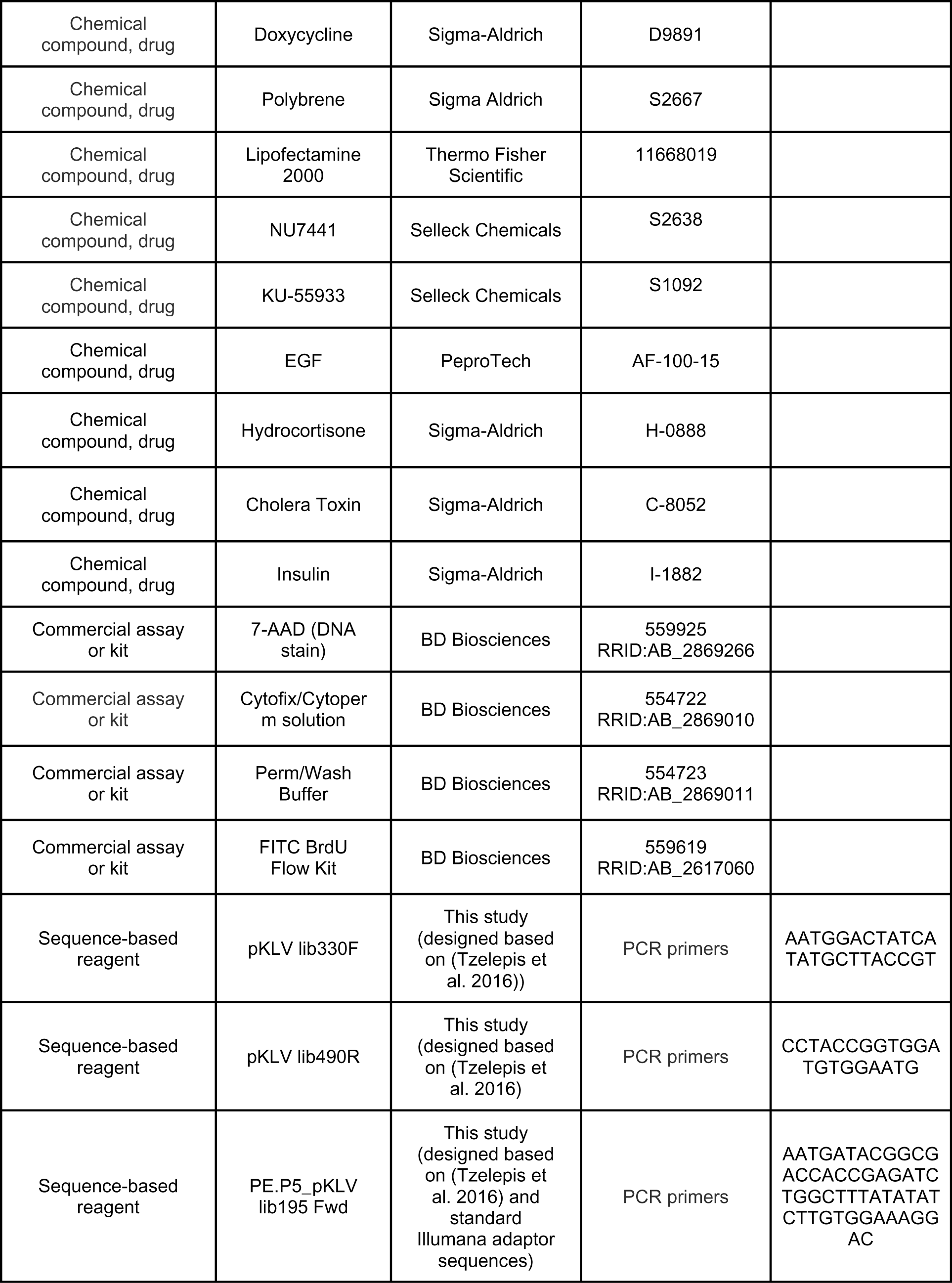
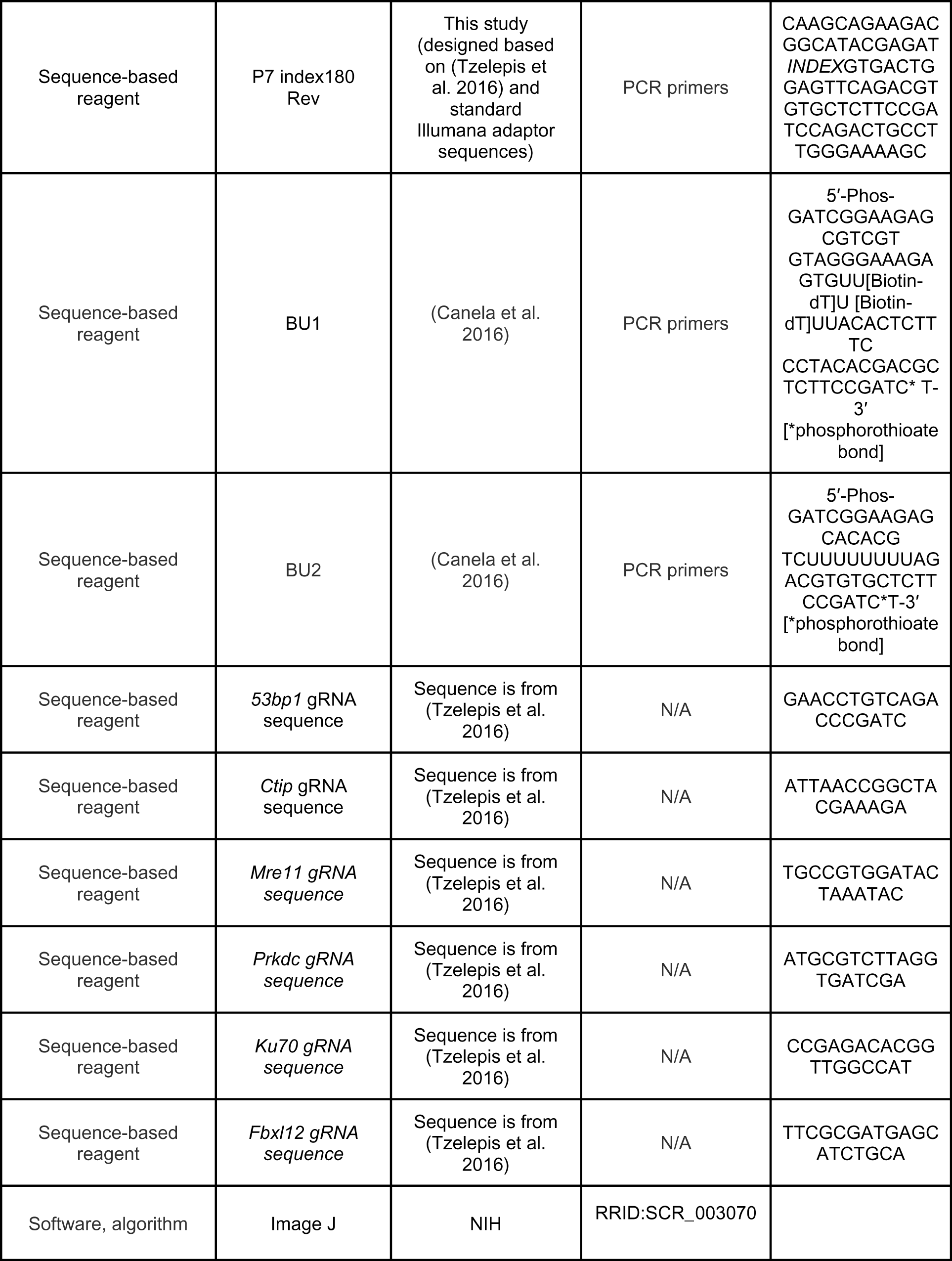
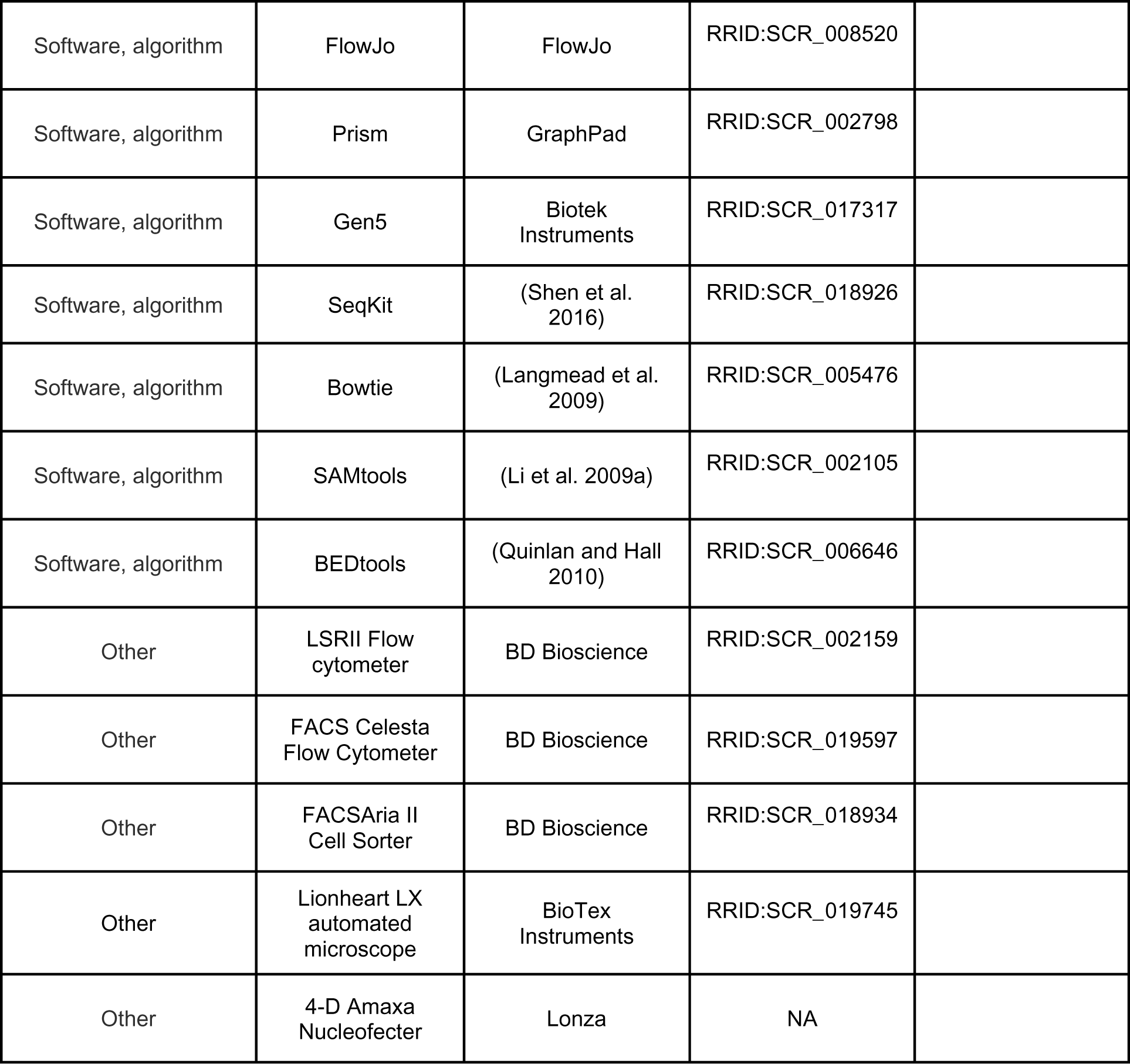

### Cell Lines and Maintenance

Abelson virus-transformed pre-B cell lines were maintained in DMEM (Thermo Fisher #11960-077) supplemented with 10% fetal bovine serum, 1% Penicillin-Streptomycin, 2 mM glutamine, 1 mM sodium pyruvate, 1X nonessential amino acids, and 0.4% beta-mercaptoethanol at 37°C with 5% CO_2_. MCF10A cells were maintained in DMEM/F12 (Gibco, #11330032), 5% horse serum, 20 ng/mL EGF, 0.5 μg/mL hydrocortisone, 100 ng/mL cholera toxin, 10 μg/mL insulin, and 1% Penicillin-Streptomycin at 37°C with 5% CO_2_. 293T cells were maintained in DMEM (Corning, #10-013-CM) supplemented with 10% fetal bovine serum and 1% Penicillin-Streptomycin at 37°C with 5% CO_2_. *Lig4^-/-^* abl pre-B cells contain pCW-Cas9 (addgene, #50661) which expresses cas9 under a doxycycline-induced promoter. To generate single cell clones of *Lig4^-/-^:53bp1^-/-^*, *Lig4^-/-^:Ku70^-/-^, Lig4^-/-^:Prkdc^-/-^,* and *Lig4^-/-^:Fbxl12^-/-^,* guide RNAs (gRNAs) against each gene were cloned into pKLV-U6gRNA-EF(BbsI)-PGKpuro2ABFP (addgene, #62348) modified to express human CD2 as a cell surface marker. *Lig4^-/-^* abl pre-B cells were grown in 3 μg/mL of doxycycline for 2 days and then nucleofected with the pKLV-gRNA plasmid using a Lonza Amaxa Nucleofector. The next day, cells were magnetically selected for human CD2 cell surface expression, and selected cells were grown in 3 μg/mL doxycycline overnight. Serial dilution in 96 well plates was used to isolate single cells. After cell growth, potential clones were confirmed to have the gene of interest knocked out by Sanger sequencing or western blotting.

### Bulk gene inactivation

gRNAs against *Mre11*, *CtIP*, and *Ku70* were cloned into pKLV-U6gRNA-EF(BbsI)-PGKpuro2ABFP (addgene, #62348). 293T cells were transfected with the pKLV-gRNA plasmid along with lentiviral packaging and lentiviral envelope plasmids. 3 days post-transfection, supernatant containing pKLV-gRNA lentivirus was filtered with a 0.45 micron filter. *Lig4^-/-^* cells were resuspended in the filtered viral supernatant supplemented with 5 μg/mL polybrene (Sigma-Aldrich, #S2667) in 6-well plates and centrifuged at 1,800 RPM for 1.5 hrs at room temperature. After spin infection, virally transduced cells were supplemented with DMEM containing 3 μg/mL doxycycline for 3 days before flow cytometry-assisted cell sorting or magnetic-assisted cell sorting based on hCD2 cell surface expression.

### Flow Cytometry

Abl pre-B cells were arrested in G_0_ using 3 μM imatinib (Selleck Chemicals, #S2475) for 48 hours. MCF10A cells were arrested in G_0_ by withdrawing EGF for 48 hours. To arrest cells in G_2_, abl pre-B cells were treated with 10 μM RO-3306 (Selleck Chemicals, #S7747) overnight. For experiments analyzing DNA-PKcs and ATM inhibition, 10 μM NU7441 (Selleck Chemicals, #S2638) or 15 μM KU-55933 (Selleck Chemicals, #S1092) was added 1 hour prior to irradiation. After irradiation with 20 Gray, cells were allowed to recover for 3 hours. Cells were then pre-extracted with 0.05% Triton-X (STI-treated abl pre-B cells), 0.2% Triton-X (proliferating abl pre-B cells) or 0.5% Triton-X (MCF10A cells) in PBS and fixed with BD Cytofix/Cytoperm solution (BD Biosciences, #554722) containing 4.2% formaldehyde. Fixed cells were stained with anti-RPA32 (Cell Signaling Technology, #2208S) for 2 hours at room temperature, and then treated with a fluorescent conjugated secondary antibody (BioLegend, #405416 or BioLegend, #405418) for 1 hour at room temperature. 7-AAD was added to each sample to stain for DNA content. Cells were analyzed using a BD LSRII Flow Cytometer or a BD FACSCelesta and flow cytometry results were further analyzed using FlowJo.

### Nuclear RPA Immunofluorescence Staining

60,000 G_0_-arrested MCF10A cells grown on cover slips were irradiated with 10 Gray and then allowed to recover for 3 hours at 37°C with 5% CO_2_. Cells were then washed with PBS containing 0.1% Tween-20 (PBST), pre-extracted using cold 0.5% Triton-X in PBS for 5 minutes, fixed with 4% formaldehyde for 15 minutes, and blocked in 3% BSA-PBST for 1 hour at room temperature. Cells were incubated overnight at 4°C in primary antibody (anti-RPA32, Cell Signaling Technology, #2208). diluted in 3% BSA-PBST Samples were then washed 3x with PBST, incubated with secondary antibody diluted in 3% BSA (Alexa Fluor 594 Goat anti-Rat IgG, BioLegend, #405422) in the dark for 1 hr at room temperature, washed 3x with PBST, and mounted in Prolong Gold Antifade Mountant with DAPI (Life Technologies, #P-36931). Images were taken using a Biotek Lionheart Automatic Microscope and foci quantification was performed using Biotek Gen5 software.

### END-Seq and RPA ChIP-Seq

Sequencing assays were performed in *Lig4^-/-^* abl pre-B cells after arrest in G_0_ with imatinib for 24 hours or arrest in G_2_ with RO-3306 for 12 hours, then treated with doxycycline for 24 hours followed by tamoxifen treatment for 4 or 8 hours to induce AsiSI breaks in the nucleus. End-seq was performed as previously described (Canela et al. 2016; Chen et al. 2021a; Wong et al. 2021). Cells were embedded in agarose plugs, lysed, and treated with proteinase K and RNase A. The DNA was then blunted with ExoVII (NEB) and ExoT (NEB), A-tailed, and ligated with a biotinylated hairpin adaptor. DNA was then recovered and sonicated to a length between 150 and 200 bp and biotinylated DNA fragments were purified using streptavidin beads (MyOne C1, Invitrogen). The DNA was then end-repaired and ligated to hairpin adaptor BU2 and amplified by PCR. RPA single-strand DNA sequencing was performed as previously described (Paiano et al. 2021). Cells were fixed in 1% formaldehyde (Sigma, F1635) for 10 min at 37°C, quenched with 125 mM glycine (Sigma), washed twice with cold 1× PBS. After centrifugation, pellets were frozen on dry ice, and stored at −80°C. Sonication, immunoprecipitation, and library preparation were performed as previously detailed (Tubbs et al. 2018). Before immunoprecipitation, sheared chromatin was precleared with 40 µL of Dynabeads Protein A (Thermo Fisher) for 30 min at 4°C. Sheared chromatin was enriched with 10 µg of anti-RPA32/RPA2 antibody (Abcam, ab10359) on Dynabeads Protein A overnight at 4°C. During library preparation, kinetic enrichment of single-strand DNA was performed by heating sheared DNA for 3 min at 95°C and allowing DNA to return to room temperature (Tubbs et al. 2018). All END-seq and RPA ChIP-seq libraries were collected by gel purification and quantified using qPCR. Sequencing was performed on the Illumina NextSeq500 (75 cycles) as previously described (Chen et al. 2021a).

### Genome Alignment and Visualization

END-seq and RPA ChIP-seq single-end reads were aligned to the mouse genome (mm10) using Bowtie v1.1.2 (Langmead et al. 2009) with parameters (-n 3 -k 1 -l 50) for END-seq and (-n 2 -m 1 -l 50) for RPA ChIP-seq. All plots or analysis were done for the top 200 *AsiSI* sites determined by END-seq. Alignment files were generated and sorted using SAMtools (Li et al. 2009b) and converted to bedgraph files using bedtools genomecov (Quinlan and Hall 2010) following by bedGraphToBigWig to make a bigwig file (Kent et al. 2010). Visualization of genomic profiles was done by the UCSC genome browser (Kent et al. 2002) and normalized to present RPM. Heat maps were produced using the R package pheatmap.

### Guide RNA Library Screen

144 million *Lig4^-/-^* abl pre-B cells were transduced with a viral tet-inducible guide RNA library (Pooled Library #67988, Addgene) containing 90,000 gRNAs targeting over 18,000 mouse genes. 3 days post-infection, cells were sorted for gRNA vector expression using a BD FACSAria flow assisted cell sorter. The next day, sorted cells were treated with 3 µg/ml doxycycline to induce gRNA expression. 7 days later, cells were treated with Gleevec to arrest cells in G_0_. 48 hours later, cells were irradiated with 20 Gray and allowed to recover for 2 hours. After collection, cells were permeabilized, fixed, and stained with anti-RPA32 in the same manner as described in the Flow Cytometry section. After staining, the top 10% and bottom 10% of RPA stained cells were collected using flow assisted cell sorting and genomic DNA was extracted. An Illumina sequencing library was generated using two rounds of PCR to amplify the gRNA and add a barcode, then purified PCR products containing the barcoded enriched gRNAs were sequenced on an Illumina HiSeq2500. Sequencing data were processed as previously described (Chen et al. 2021a).

### Western Blotting

The following antibodies were used for western blot analysis: CtIP (gift from Dr. Richard Baer, [Columbia University, New York], 1:1000), MRE11 (Novus Biologicals, NB100-142, 1:2000), GAPDH (Sigma, G8795, 1:10,000), DNA-PK (Invitrogen, MA5-32192, 1:1000), KAP1 (Genetex, GTX102226, 1:2000), KU70 (Cell Signaling Technology, #4588, 1:1000).

### Plasmid Constructs

pCW-Cas9 was a gift from Eric Lander and David Sabatini (Addgene plasmid #50661) (Wang et al. 2014). pKLV-U6gRNA(BbsI)-PGKpuro2ABFP was a gift from Kosuke Yusa (Addgene plasmid #50946) (Koike-Yusa et al. 2014). Mouse Improved Genome-wide Knockout CRISPR Library v2 was a gift from Kosuke Yusa (Addgene #67988) (Tzelepis et al. 2016).

## Notes

### Competing Interest Statement

The authors have declared no competing interest.

## References

Averbeck NB, Ringel O, Herrlitz M, Jakob B, Durante M, Taucher-Scholz G. 2014. DNA end resection is needed for the repair of complex lesions in G_1_-phase human cells. Cell Cycle 13: 2509–2516.

Barlow JH, Lisby M, Rothstein R. 2008. Differential regulation of the cellular response to DNA double-strand breaks in G_1_. Mol Cell 30: 73–85.

Bartlett EJ, Lees-Miller SP. 2018. Established and Emerging Roles of the DNA-Dependent Protein Kinase Catalytic Subunit (DNA-PKcs). in Targeting the DNA Damage Response for Anti-Cancer Therapy (eds. J Pollard, N Curtin), pp. 315–338. Springer International Publishing, Cham.

Biehs R, Steinlage M, Barton O, Juhász S, Künzel J, Spies J, Shibata A, Jeggo PA, Löbrich M. 2017. DNA Double-Strand Break Resection Occurs during Non-homologous End Joining in G_1_ but Is Distinct from Resection during Homologous Recombination. Mol Cell 65: 671–684.e675.

Blackford AN, Jackson SP. 2017. ATM, ATR, and DNA-PK: The Trinity at the Heart of the DNA Damage Response. Mol Cell 66: 801–817.

Bredemeyer AL, Sharma GG, Huang CY, Helmink BA, Walker LM, Khor KC, Nuskey B, Sullivan KE, Pandita TK, Bassing CH et al. 2006. ATM stabilizes DNA double-strand-break complexes during V(D)J recombination. Nature 442: 466–470.

Britton S, Chanut P, Delteil C, Barboule N, Frit P, Calsou P. 2020. ATM antagonizes NHEJ proteins assembly and DNA-ends synapsis at single-ended DNA double strand breaks. Nucleic Acids Res 48: 9710–9723.

Bunting SF, Callén E, Wong N, Chen HT, Polato F, Gunn A, Bothmer A, Feldhahn N, Fernandez-Capetillo O, Cao L et al. 2010. 53BP1 inhibits homologous recombination in Brca1-deficient cells by blocking resection of DNA breaks. Cell 141: 243–254.

Canela A, Sridharan S, Sciascia N, Tubbs A, Meltzer P, Sleckman BP, Nussenzweig A. 2016. DNA Breaks and End Resection Measured Genome-wide by End Sequencing. Mol Cell 63: 898–911.

Chaplin AK, Hardwick SW, Liang S, Kefala Stavridi A, Hnizda A, Cooper LR, De Oliveira TM, Chirgadze DY, Blundell TL. 2021. Dimers of DNA-PK create a stage for DNA double-strand break repair. Nat Struct Mol Biol 28: 13–19.

Chen BR, Wang Y, Tubbs A, Zong D, Fowler FC, Zolnerowich N, Wu W, Bennett A, Chen CC, Feng W et al. 2021a. LIN37-DREAM prevents DNA end resection and homologous recombination at DNA double-strand breaks in quiescent cells. Elife 10.

Chen X, Xu X, Chen Y, Cheung JC, Wang H, Jiang J, de Val N, Fox T, Gellert M, Yang W. 2021b. Structure of an activated DNA-PK and its implications for NHEJ. Mol Cell 81: 801–810.e803.

Clerici M, Mantiero D, Guerini I, Lucchini G, Longhese MP. 2008. The Yku70-Yku80 complex contributes to regulate double-strand break processing and checkpoint activation during the cell cycle. EMBO Rep 9: 810–818.

Dev H, Chiang TW, Lescale C, de Krijger I, Martin AG, Pilger D, Coates J, Sczaniecka-Clift M, Wei W, Ostermaier M et al. 2018. Shieldin complex promotes DNA end-joining and counters homologous recombination in BRCA1-null cells. Nat Cell Biol 20: 954–965.

Feng L, Chen J. 2012. The E3 ligase RNF8 regulates KU80 removal and NHEJ repair. Nat Struct Mol Biol 19: 201–206.

Forment JV, Walker RV, Jackson SP. 2012. A high-throughput, flow cytometry-based method to quantify DNA-end resection in mammalian cells. Cytometry A 81: 922–928.

Golub EI, Gupta RC, Haaf T, Wold MS, Radding CM. 1998. Interaction of human rad51 recombination protein with single-stranded DNA binding protein, RPA. Nucleic Acids Res 26: 5388–5393.

Gottlieb TM, Jackson SP. 1993. The DNA-dependent protein kinase: requirement for DNA ends and association with Ku antigen. Cell 72: 131–142.

Gravel S, Chapman JR, Magill C, Jackson SP. 2008. DNA helicases Sgs1 and BLM promote DNA double-strand break resection. Genes Dev 22: 2767–2772.

Hammarsten O, Chu G. 1998. DNA-dependent protein kinase: DNA binding and activation in the absence of Ku. Proc Natl Acad Sci U S A 95: 525–530.

Ishida N, Nakagawa T, Iemura SI, Yasui A, Shima H, Katoh Y, Nagasawa Y, Natsume T, Igarashi K, Nakayama K. 2017. Ubiquitylation of Ku80 by RNF126 Promotes Completion of Nonhomologous End Joining-Mediated DNA Repair. Mol Cell Biol 37.

Ismail IH, Gagné JP, Genois MM, Strickfaden H, McDonald D, Xu Z, Poirier GG, Masson JY, Hendzel MJ. 2015. The RNF138 E3 ligase displaces Ku to promote DNA end resection and regulate DNA repair pathway choice. Nat Cell Biol 17: 1446–1457.

Jackson SP, Bartek J. 2009. The DNA-damage response in human biology and disease. Nature 461: 1071–1078.

Kent WJ, Sugnet CW, Furey TS, Roskin KM, Pringle TH, Zahler AM, Haussler D. 2002. The human genome browser at UCSC. Genome Res 12: 996–1006.

Kent WJ, Zweig AS, Barber G, Hinrichs AS, Karolchik D. 2010. BigWig and BigBed: enabling browsing of large distributed datasets. Bioinformatics 26: 2204–2207.

Koike-Yusa H, Li Y, Tan EP, Velasco-Herrera MeC, Yusa K. 2014. Genome-wide recessive genetic screening in mammalian cells with a lentiviral CRISPR-guide RNA library. Nat Biotechnol 32: 267–273.

Langmead B, Trapnell C, Pop M, Salzberg SL. 2009. Ultrafast and memory-efficient alignment of short DNA sequences to the human genome. Genome Biol 10: R25.

Lee SE, Moore JK, Holmes A, Umezu K, Kolodner RD, Haber JE. 1998. Saccharomyces Ku70, mre11/rad50 and RPA proteins regulate adaptation to G_2_/M arrest after DNA damage. Cell 94: 399-409.

Li H, Handsaker B, Wysoker A, Fennell T, Ruan J, Homer N, Marth G, Abecasis G, Durbin R, Genome Project Data Processing S. 2009a. The Sequence Alignment/Map format and SAMtools. Bioinformatics 25: 2078-2079.

Li H, Handsaker B, Wysoker A, Fennell T, Ruan J, Homer N, Marth G, Abecasis G, Durbin R, Subgroup GPDP. 2009b. The Sequence Alignment/Map format and SAMtools. Bioinformatics 25: 2078–2079.

Ma Y, Pannicke U, Schwarz K, Lieber MR. 2002. Hairpin opening and overhang processing by an Artemis/DNA-dependent protein kinase complex in nonhomologous end joining and V(D)J recombination. Cell 108: 781–794.

Ma Y, Schwarz K, Lieber MR. 2005. The Artemis:DNA-PKcs endonuclease cleaves DNA loops, flaps, and gaps. DNA Repair (Amst) 4: 845–851.

Menon V, Povirk LF. 2016. End-processing nucleases and phosphodiesterases: An elite supporting cast for the non-homologous end joining pathway of DNA double-strand break repair. DNA Repair (Amst) 43: 57–68.

Mimitou EP, Symington LS. 2008. Sae2, Exo1 and Sgs1 collaborate in DNA double-strand break processing. Nature 455: 770-774.

Mirman Z, Lottersberger F, Takai H, Kibe T, Gong Y, Takai K, Bianchi A, Zimmermann M, Durocher D, de Lange T. 2018. 53BP1-RIF1-shieldin counteracts DSB resection through CST- and Polα-dependent fill-in. Nature 560: 112–116.

Noordermeer SM, Adam S, Setiaputra D, Barazas M, Pettitt SJ, Ling AK, Olivieri M, Álvarez-Quilón A, Moatti N, Zimmermann M et al. 2018. The shieldin complex mediates 53BP1-dependent DNA repair. Nature 560: 117–121.

Paiano J, Zolnerowich N, Wu W, Pavani R, Wang C, Li H, Zheng L, Shen B, Sleckman BP, Chen BR et al. 2021. Role of 53BP1 in end protection and DNA synthesis at DNA breaks. Genes Dev 35: 1356–1367.

Paull TT, Gellert M. 1998. The 3’ to 5’ exonuclease activity of Mre 11 facilitates repair of DNA double-strand breaks. Mol Cell 1: 969–979.

Postow L, Funabiki H. 2013. An SCF complex containing Fbxl12 mediates DNA damage-induced Ku80 ubiquitylation. Cell Cycle 12: 587–595.

Postow L, Ghenoiu C, Woo EM, Krutchinsky AN, Chait BT, Funabiki H. 2008. Ku80 removal from DNA through double strand break-induced ubiquitylation. J Cell Biol 182: 467–479.

Quinlan AR, Hall IM. 2010. BEDTools: a flexible suite of utilities for comparing genomic features. Bioinformatics 26: 841–842.

San Filippo J, Sung P, Klein H. 2008. Mechanism of eukaryotic homologous recombination. Annu Rev Biochem 77: 229–257.

Sartori AA, Lukas C, Coates J, Mistrik M, Fu S, Bartek J, Baer R, Lukas J, Jackson SP. 2007. Human CtIP promotes DNA end resection. Nature 450: 509–514.

Scully R, Panday A, Elango R, Willis NA. 2019. DNA double-strand break repair-pathway choice in somatic mammalian cells. Nat Rev Mol Cell Biol 20: 698–714.

Setiaputra D, Durocher D. 2019. Shieldin - the protector of DNA ends. EMBO Rep 20.

Shao Z, Davis AJ, Fattah KR, So S, Sun J, Lee KJ, Harrison L, Yang J, Chen DJ. 2012. Persistently bound Ku at DNA ends attenuates DNA end resection and homologous recombination. DNA Repair (Amst) 11: 310–316.

Shen W, Le S, Li Y, Hu F. 2016. SeqKit: A Cross-Platform and Ultrafast Toolkit for FASTA/Q File Manipulation. PLoS One 11: e0163962.

Stiff T, O’Driscoll M, Rief N, Iwabuchi K, Löbrich M, Jeggo PA. 2004. ATM and DNA-PK function redundantly to phosphorylate H2AX after exposure to ionizing radiation. Cancer Res 64: 2390–2396.

Sugiyama T, Kowalczykowski SC. 2002. Rad52 protein associates with replication protein A (RPA)-single-stranded DNA to accelerate Rad51-mediated displacement of RPA and presynaptic complex formation. J Biol Chem 277: 31663–31672.

Symington LS, Gautier J. 2011. Double-strand break end resection and repair pathway choice. Annu Rev Genet 45: 247–271.

Tkáč J, Xu G, Adhikary H, Young JTF, Gallo D, Escribano-Díaz C, Krietsch J, Orthwein A, Munro M, Sol W et al. 2016. HELB Is a Feedback Inhibitor of DNA End Resection. Mol Cell 61: 405–418.

Trujillo KM, Yuan SS, Lee EY, Sung P. 1998. Nuclease activities in a complex of human recombination and DNA repair factors Rad50, Mre11, and p95. J Biol Chem 273: 21447-21450.

Tubbs A, Sridharan S, van Wietmarschen N, Maman Y, Callen E, Stanlie A, Wu W, Wu X, Day A, Wong N et al. 2018. Dual Roles of Poly(dA:dT) Tracts in Replication Initiation and Fork Collapse. Cell 174: 1127–1142.e1119.

Tzelepis K, Koike-Yusa H, De Braekeleer E, Li Y, Metzakopian E, Dovey OM, Mupo A, Grinkevich V, Li M, Mazan M et al. 2016. A CRISPR Dropout Screen Identifies Genetic Vulnerabilities and Therapeutic Targets in Acute Myeloid Leukemia. Cell Rep 17: 1193–1205.

Wang H, Shi LZ, Wong CC, Han X, Hwang PY, Truong LN, Zhu Q, Shao Z, Chen DJ, Berns MW et al. 2013. The interaction of CtIP and Nbs1 connects CDK and ATM to regulate HR-mediated double-strand break repair. PLoS Genet 9: e1003277.

Wang T, Wei JJ, Sabatini DM, Lander ES. 2014. Genetic screens in human cells using the CRISPR-Cas9 system. Science 343: 80–84.

Weinfeld M, Soderlind KJ. 1991. 32P-postlabeling detection of radiation-induced DNA damage: identification and estimation of thymine glycols and phosphoglycolate termini. Biochemistry 30: 1091–1097.

Wong N, John S, Nussenzweig A, Canela A. 2021. END-seq: An Unbiased, High-Resolution, and Genome-Wide Approach to Map DNA Double-Strand Breaks and Resection in Human Cells. Methods Mol Biol 2153: 9–31.

Wright WD, Shah SS, Heyer WD. 2018. Homologous recombination and the repair of DNA double-strand breaks. J Biol Chem 293: 10524–10535.

Yaneva M, Kowalewski T, Lieber MR. 1997. Interaction of DNA-dependent protein kinase with DNA and with Ku: biochemical and atomic-force microscopy studies. EMBO J 16: 5098–5112.

Zahid S, Seif El Dahan M, Iehl F, Fernandez-Varela P, Le Du MH, Ropars V, Charbonnier JB. 2021. The Multifaceted Roles of Ku70/80. Int J Mol Sci 22.

Zha S, Jiang W, Fujiwara Y, Patel H, Goff PH, Brush JW, Dubois RL, Alt FW. 2011. Ataxia telangiectasia-mutated protein and DNA-dependent protein kinase have complementary V(D)J recombination functions. Proc Natl Acad Sci U S A 108: 2028–2033.

Zha S, Shao Z, Zhu Y. 2021. The plié by DNA-PK: dancing on DNA. Mol Cell 81: 644–646.

Zhou Y, Paull TT. 2013. DNA-dependent protein kinase regulates DNA end resection in concert with Mre11-Rad50-Nbs1 (MRN) and ataxia telangiectasia-mutated (ATM). J Biol Chem 288: 37112–37125.

Zhu Z, Chung WH, Shim EY, Lee SE, Ira G. 2008. Sgs1 helicase and two nucleases Dna2 and Exo1 resect DNA double-strand break ends. Cell 134: 981–994.

